# Tumor suppressor-like mutations mark a dosage-constrained dependency in leukemia

**DOI:** 10.64898/2026.09.29.755484

**Authors:** Jonathan D. Lee, Madison M. Uyemura, Francisca Lamini, Ryan R. Posey, Ellie H. Shin, Amyah C. Harris, Tiziana Ottone, Giulia Cheloni, David B. Chou, Bon Q. Trinh, Joao A. Paulo, Daniel G. Tenen, John G. Clohessy, Maria Teresa Voso, Timothy A. Graubert, Frank J. Slack

## Abstract

Cancer genes are classically categorized as oncogenes or tumor suppressors based on whether gain or loss of function promotes malignancy. Here we show that oncogene-imposed dosage constraints can produce mutations that mimic tumor suppressor loss yet mark essential genes. In acute promyelocytic leukemia (APL), the transcription factor *EBF3* gene harbors recurrent heterozygous loss-of-function mutations, a tumor suppressor signature, and is essential for APL survival. We resolve this paradox by showing that while PML::RARA, the fusion oncoprotein defining APL, drives EBF3 expression, excess EBF3 suppresses proliferation. Patient mutations reduce EBF3 activity through heterozygous inactivation, optimizing its expression within a corridor bounded by fitness constraints at both extremes. PML::RARA activates *EBF3* via a p300-dependent intronic enhancer, and p300 inhibition exploits this constraint by suppressing EBF3 below the viability threshold. Our findings reveal that oncogenes can impose dosage constraints that drive adaptive mutations conventionally misclassified as tumor suppressor loss, with implications for interpreting cancer genomes.

## Introduction

Cancer genes are typically classified as oncogenes, which promote malignancy when activated, or tumor suppressors, which restrain it until lost^1–4^. This binary framework shapes how mutations are interpreted: gain-of-function implicates oncogenes, loss-of-function implicates tumor suppressors. Context-dependent exceptions exist, such as the same gene acting as an oncogene or a tumor suppressor, but these occur across distinct disease settings^5–8^. A gene exhibiting tumor suppressor mutation patterns (heterozygous, loss-of-function, clustered in functional domains) while being essential for survival within the same cancer context would challenge this interpretive framework. Whether such mutations occur in human cancer has not been established.

Acute promyelocytic leukemia (APL) is defined by the t(15;17) translocation, which generates the PML::RARA fusion oncoprotein^9–12^. PML::RARA was long considered a transcriptional repressor that blocks myeloid differentiation by silencing retinoic acid target genes^13–16^. Recent studies, however, have revealed a more complex picture: PML::RARA exerts both repressive and activating functions, the latter through recruitment of p300 and formation of super-enhancers^17–19^. These activating functions appear critical for leukemogenesis, and PML::RARA-activated target genes such as *GFI1* are necessary for APL cell maintenance^18^. However, many target genes including *GFI1* are broadly required across myeloid malignancies, and the dependencies that uniquely define APL remain unclear^20,21^.

A closer examination of APL and AML genome sequencing studies reveals recurrent somatic mutations in *EBF3* in APL but not other myeloid malignancies^22–25^. EBF3 is a helix-loop-helix transcription factor with essential roles in neuronal development^26^. Germline heterozygous mutations cause HADDS, a neurodevelopmental disorder, establishing *EBF3* as haploinsufficient^27–30^. In cancers including acute myeloid leukemia, EBF3 has been characterized as a tumor suppressor, with its ectopic expression inducing cell cycle arrest and apoptosis^31–34^. Strikingly, APL mutations mirror HADDS: heterozygous, clustered in the DNA-binding domain, and predicted loss-of-function. Although this pattern is consistent with tumor suppression, the APL-specific occurrence and absence of homozygous loss suggest a more complex constraint.

Here we report that EBF3, despite its reported tumor suppressor activity in other contexts, is highly expressed in APL and essential for leukemic cell survival. *EBF3* knockdown phenocopies PML::RARA loss, establishing EBF3 as a major effector of the fusion’s transcriptional program. Unexpectedly, EBF3 is constrained at both extremes: loss triggers apoptosis, while excess suppresses growth. Patient-derived *EBF3* mutations are heterozygous, with the wild-type allele expressed at normal levels, revealing selection for reduced but not absent activity. PML::RARA maintains *EBF3* within this constrained range via a p300-dependent intronic enhancer, and p300 inhibition exploits this dependency by suppressing *EBF3* below the viability threshold. Our findings reveal that fusion oncogenes can impose dosage constraints that shape tumor evolution and create therapeutically actionable vulnerabilities.

## Results

### EBF3 is an APL-specific dependency downstream of PML::RARA

APL is transcriptionally and histopathologically distinct from other myeloid malignancies^23^. The PML::RARA fusion defines and maintains the APL state, and its degradation or depletion via ATRA (all-trans retinoic acid), ATO (arsenic trioxide), or genetic knockdown collapses this state and eliminates leukemic cells^35–37^ (Fig. 1A). Prior studies have identified transcription factors contributing to the APL state through transcriptomic and chromatin-based approaches^18,38,39^. However, many of these transcription factors have established roles across other leukemias. Whether any transcription factors downstream of PML::RARA uniquely support the APL state remains unclear.

**Figure 1.**
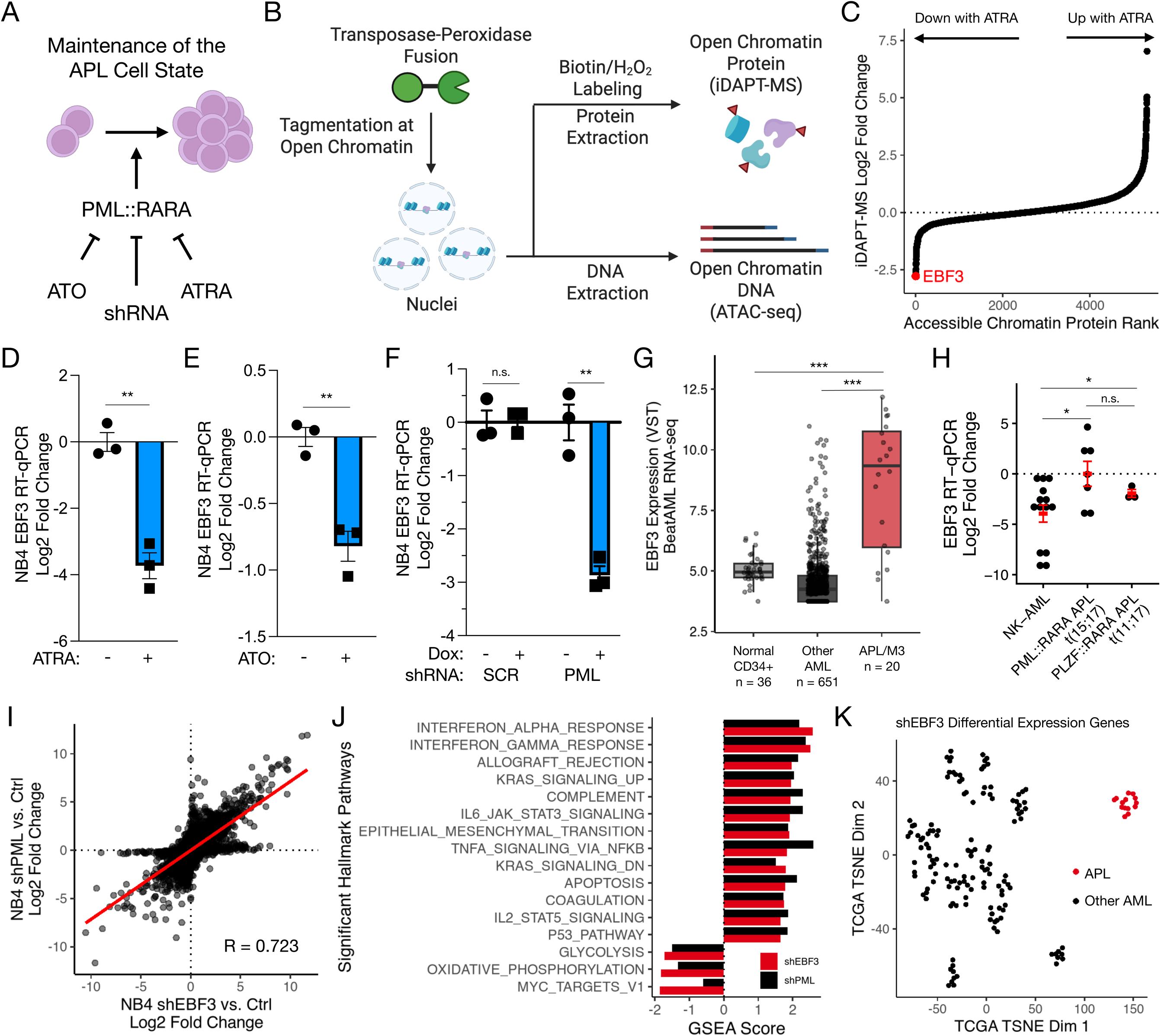
EBF3 is an APL-specific, PML::RARA-dependent transcription factor. (A) Schematic of APL cell state maintenance via PML::RARA and modes of suppression. (B) Schematic of the iDAPT-MS^45^ (integrative DNA And Protein Tagging with mass spectrometry) approach. A recombinant transposase-peroxidase fusion protein is added to nuclei, in which the transposase domain binds to and tags DNA at open chromatin. The peroxidase domain catalyzes transposase-proximal biotin labeling upon hydrogen peroxide addition, thereby tagging open chromatin-associated proteins for mass spectrometry analysis (iDAPT-MS). DNA may also be extracted for parallel ATAC-seq analysis. (C) NB4 cells were treated with 1 μM ATRA or DMSO for 48 hours, and transcription factors at accessible chromatin were identified by iDAPT-MS^45^. EBF3 was the top factor lost upon ATRA-induced treatment (average log2 fold change rank plot). n = 3/condition. (D) *EBF3* RNA expression (RT-qPCR) of NB4 cells following 1 μM ATRA or DMSO treatment for 48 hours. *ACTB* RNA was used as an internal control. n = 3. **, p < 0.01. Welch’s t-test. (E) *EBF3* RNA expression (RT-qPCR) of NB4 cells following 1 μM ATO or vehicle treatment for 48 hours. *ACTB* RNA was used as an internal control. n = 3. **, p < 0.01. Welch’s t-test. (F) *EBF3* RNA expression (RT-qPCR) of NB4 cells harboring doxycycline-inducible shRNA transgenes (SCR or PML; SCR, scrambled control) and following 1 μg/mL doxycycline or vehicle treatment for 72 hours. *ACTB* RNA was used as an internal control. n = 3. n.s., not significant. **, p < 0.01. Welch’s t-test. (G) *EBF3* RNA expression (RNA-seq) of APL or FAB M3 patient samples compared to normal CD34+/bone marrow hematopoietic progenitors and other myeloid malignancies (AML). Data from the BeatAML cohort^49^. ***, p < 0.001. Wilcoxon rank-sum test. (H) *EBF3* RNA expression (RT-qPCR) of PML::RARA t(15;17) (n = 7) and PLZF::RARA t(11;17) (n = 3) APL patient samples compared to normal karyotype AML (NK-AML; n = 14). *ACTB* RNA was used as an internal control. *, p < 0.05. Welch’s t-test. (I) Distribution of gene expression changes (RNA-seq) upon *EBF3* knockdown versus PML::RARA knockdown in NB4 cells (Pearson’s R = 0.723). (J) MSigDB^83^ Hallmark Pathway gene set enrichment analysis (GSEA) of genes dysregulated upon EBF3 or PML::RARA shRNA knockdown, relative to negative controls (scrambled shRNA or empty vector). Shown pathways are those with FDR < 0.01 upon *EBF3* knockdown. GSEA score, normalized enrichment score from fgsea. (K) t-SNE projection of TCGA-LAML^23^ RNA-seq profiles using significant differentially expressed genes (FDR < 0.05, |Log2 Fold Change| > 1) upon shEBF3 knockdown as features.

Most transcription factors regulate gene expression through binding at accessible chromatin^40,41^. However, standard motif-based discovery approaches cannot directly identify chromatin-bound proteins and may miss factors with degenerate or context-dependent binding^42–44^. To overcome this limitation, we previously developed iDAPT-MS^45,46^ (integrative DNA And Protein Tagging with mass spectrometry) to profile open chromatin-associated proteins in NB4 cells and their dynamics upon ATRA-induced differentiation (Fig. 1B). Examining this dataset for individual transcription factor changes, we identified EBF3 as the most significantly depleted protein at accessible chromatin after 48 hours of ATRA treatment (Fig. 1C, Supplemental Fig. 1A). To validate EBF3 endogenous protein expression, we engineered NB4 cells harboring a C-terminal HiBiT-3xFLAG tag at the *EBF3* gene via CRISPR/Cas9 (Supplemental Fig. 1B-D). Nuclear/cytoplasmic fractionation showed that EBF3 is strongly enriched in the nucleus (Supplemental Fig. 1E). From fractionation analysis, we found EBF3 protein in the nucleus is decreased upon ATRA, ATO, or PML-targeting shRNA (Supplemental Fig. 1F-I). EBF3 decreased upon ATRA treatment at both protein and RNA levels; ATO and PML-targeting shRNA also reduced *EBF3* RNA, supporting transcriptional regulation via PML::RARA (Fig. 1D-F, Supplemental Fig. 1J). Critically, ATRA reduced *EBF3* RNA expression in primary APL samples *ex vivo* and in patients receiving therapy, consistent with EBF3’s transcriptional regulation by PML::RARA in patients (Supplemental Fig. 1K-L).

We next examined whether EBF3 expression is specific to PML::RARA and the APL state. Across 217 hematopoietic cancer cell lines profiled in the Cancer Cell Line Encyclopedia^47,48^, the NB4 cell line, the only APL cell line in this group, exhibits the second highest *EBF3* expression, 4.88 standard deviations above the mean (Supplemental Fig. 2A). In both BeatAML^22,49^ and TCGA-LAML^23^ cohorts, *EBF3* was highly expressed in APL compared to other myeloid malignancies, a finding we validated in an independent in-house cohort (Fig. 1G-H, Supplemental Fig. 2B). Compared to normal CD34+ and bone marrow mononuclear cells in the BeatAML cohort^49^ and normal promyelocytes in the TCGA cohort^50^, *EBF3* is highly expressed in APL samples (Fig. 1G, Supplemental Fig. 2C).

*EBF3* expression alone distinguished APL from non-APL AML with high accuracy (BeatAML AUC = 0.91, TCGA-LAML AUC = 0.95; Supplemental Fig. 2D-E), though it does not stratify patients by Sanz clinical risk^24,51^, suggesting *EBF3* marks APL identity rather than disease severity (Supplemental Fig. 2F). Notably, *EBF3* was not among the top-ranked differentially expressed genes by fold-change (ranking #25 in BeatAML and #210 in TCGA-LAML; Supplemental Fig. 2G-I), illustrating that proteomic discovery via iDAPT-MS can identify functionally important factors that transcriptomic approaches might deprioritize. For subsequent experiments, we used RNA expression as a practical readout of EBF3 levels.

Beyond PML::RARA APL, we found that *EBF3* expression is sustained in APL-like disease driven by PLZF::RARA t(11;17)^14,52^, FIP1L1::RARA t(4;17)^53,54^, and STRN3::RARA t(14;17)^55^ fusions (Fig. 1H, Supplemental Fig. 2J-K). These rare APL subtypes are often resistant to ATRA, ATO, or both^56^; the conservation of EBF3 expression across these fusions raises the possibility that *EBF3* represents a shared vulnerability.

Given EBF3’s role as a transcription factor, these data suggested that PML::RARA’s transcriptional program is at least partially driven by EBF3. To test this, we compared gene expression changes upon shRNA-mediated *EBF3* knockdown versus PML::RARA knockdown relative to negative shRNA controls by RNA-seq (Fig. 1I, Supplemental Fig. 2L-M). Transcriptional changes upon *PML::RARA* and *EBF3* knockdown were highly correlated (Pearson’s R = 0.72 among genes with FDR < 0.05 in either contrast), and pathway analysis of the transcriptional changes showed substantial overlap between both perturbations (Fig. 1I-J). To assess whether the *EBF3* gene expression program uniquely defines the APL state in human patients, we performed t-SNE analysis of BeatAML^49^ and TCGA-LAML^23^ RNA-seq cohorts using only significant differentially expressed genes upon *EBF3* knockdown (FDR < 0.05, |log2 fold change| > 1; 1,559 genes). We found that in both cohorts, APL samples formed a distinct, separate cluster removed from other AML samples (Fig. 1K, Supplemental Fig. 2N). These findings substantiate EBF3’s critical role in mediating the APL transcriptional state.

### EBF3 is essential for APL survival

We next sought to test whether EBF3 is functionally required for APL cell survival. Analysis of DepMap CRISPR/Cas9-based screening^57,58^ data revealed that NB4, the only APL cell line in the dataset, is the sole hematopoietic cancer cell line with *EBF3* dependency (Fig. 2A). In contrast, the PML::RARA target gene *GFI1* is broadly essential across hematopoietic cancer cell lines^18^ (Supplemental Fig. 3A). Among the top 10 differentially abundant TFs identified by iDAPT-MS, only *EBF3* and *RARA* (of PML::RARA fusion) showed dependency in NB4 (Supplemental Fig. 3B).

**Figure 2.**
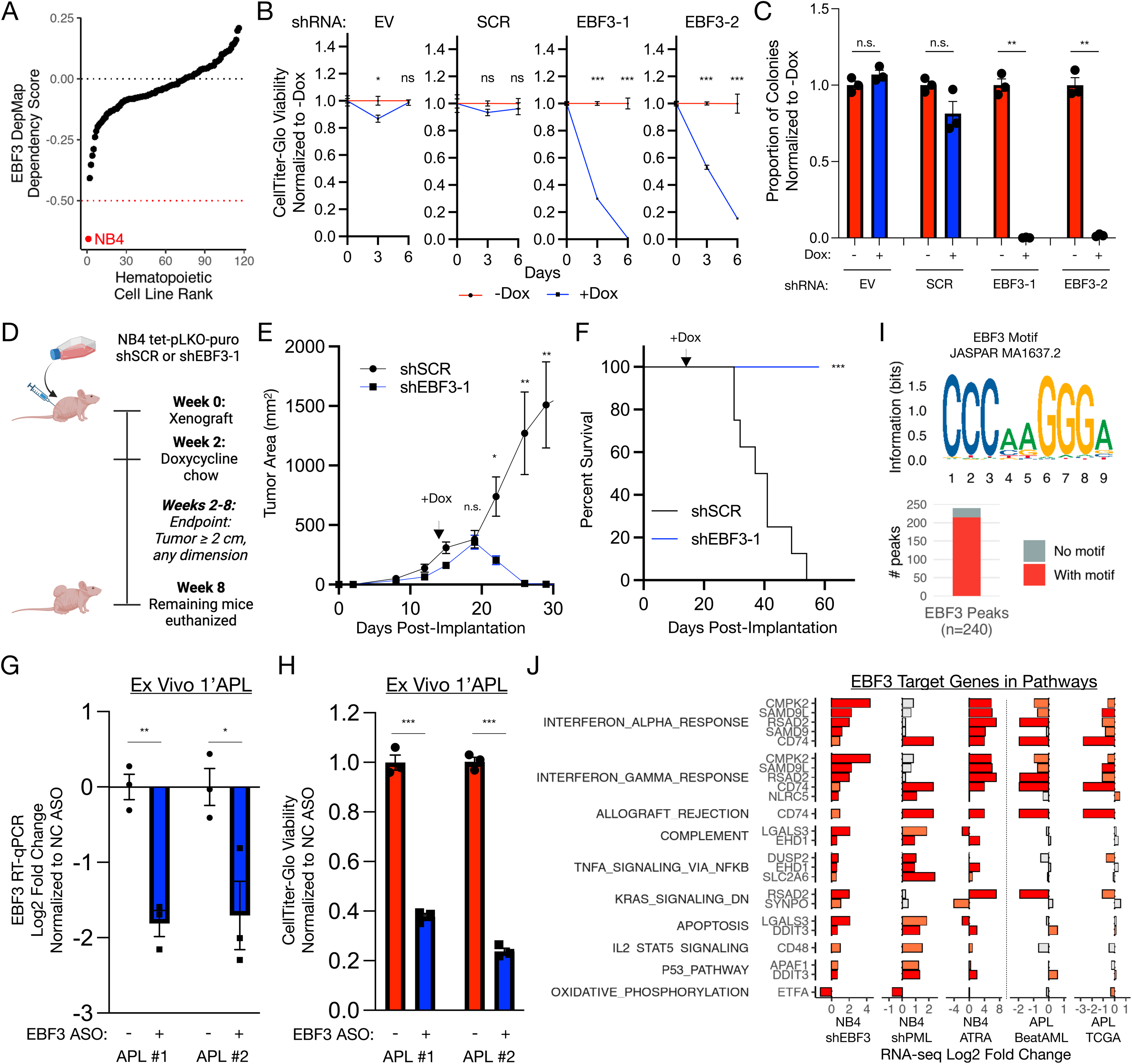
EBF3 is essential for APL cell survival. (A) EBF3 DepMap^57,58^ 23Q4 Chronos dependency scores across hematopoietic cancer cell lines (CRISPR/Cas9 knockout-based). NB4, the only APL cell line, is the only cell line with strong EBF3 dependency (Chronos score < -0.5). (B) CellTiter-Glo viability of NB4 cells harboring doxycycline-inducible *EBF3* or negative control shRNA transgenes (SCR, scrambled; EV, empty vector) and treated with or without 1 μg/mL doxycycline for the indicated time. Luminescence is normalized to the corresponding -Dox condition. n = 5. ns, not significant. *, p < 0.05. ***, p < 0.001. Welch’s t-test. (C) Colony formation unit (CFU) assay of NB4 cells harboring doxycycline-inducible *EBF3* or negative control shRNA transgenes and treated with or without 1 μg/mL doxycycline for 14 days. n = 3. n.s., not significant. **, p < 0.01. Welch’s t-test. (D) Schematic of mouse xenograft experiment. After subcutaneous injection of NB4 cells harboring doxycycline-inducible shRNAs (SCR or EBF3-1), tumors were allowed to engraft without perturbation. At two weeks when tumors were palpable, mice were placed on doxycycline chow. Tumor size was monitored, and endpoint was determined when tumor size reached 2 cm or more along any dimension. (E) Tumor area (mm^2^) over time in mice xenografted with NB4 cells transduced with SCR or *EBF3* shRNA. n = 10/group. n.s., not significant. *, p < 0.05. **, p < 0.01. Welch’s t-test. (F) Percent survival over time of mouse xenograft experiment. n = 10/group. ***, p < 0.001. Log-rank test. (G) *EBF3* RNA expression (RT-qPCR) of primary APL patient samples upon EBF3 (+) or nontargeting (-) ASO treatment. *ACTB* RNA was used as an internal control. n = 3. *, p < 0.05. **, p < 0.01. Welch’s t-test. (H) CellTiter-Glo viability of primary APL patient samples following EBF3 (+) or nontargeting (-) ASO treatment. Luminescence is normalized to the corresponding nontargeting ASO condition. n = 3. ***, p < 0.001. Welch’s t-test. (I) Top, EBF3 motif logo used for motif enrichment (JASPAR^84^ MA1637.2). Bottom, EBF3 CUT&RUN peaks (IDR [irreproducible discovery rate] < 5%) with or without canonical EBF3 motif enrichment (n = 240 peaks, 90% with motif). (J) Differential gene expression (log2 fold change) of EBF3 direct target genes in pathways defined in Figure 1H across multiple comparisons. Red, significant at the FDR < 0.01 level; light red, significant at the FDR < 0.05 level. Gray, not significant.

To validate this finding, we knocked down *EBF3* using two independent doxycycline-inducible shRNAs, with scrambled shRNA and empty vector as negative controls (Supplemental Fig. 2L-M). *EBF3* knockdown reduced viability and eliminated clonogenic capacity (Fig. 2B-C). Similar results were obtained with the NB4-R2 cell line, an ATRA-resistant subclone of NB4^59^ (Supplemental Fig. 3C-D). In murine xenograft models, *EBF3* knockdown after tumor engraftment significantly reduced tumor burden and extended survival (Fig. 2D-F). To confirm these findings in patient samples, we knocked down *EBF3* in primary APL cells using antisense oligonucleotides (ASOs). *EBF3* knockdown reduced viability in both patient samples tested (Fig. 2G-H). Together, these data establish that *EBF3* is essential for APL survival in cell lines, xenografts, and primary patient samples.

To understand how EBF3 loss modifies viability and transcription (Fig. 1I-J, 2A-B), we performed CUT&RUN^60^ to identify direct EBF3 targets. EBF3 bound a restricted set of 240 genomic loci, 90% containing the canonical EBF motif (Fig. 2I). *EBF3* knockdown increased chromatin accessibility at its binding sites and moderately de-repressed proximal genes (407 target genes) (Supplemental Fig. 3E-F). Direct targets upregulated upon *EBF3* knockdown included the interferon response genes *CD74, CMPK2, RSAD2,* and *SAMD9/SAMD9L*, and an established driver of apoptosis, *DDIT3* (Fig. 2J, Supplemental Fig. 3G-J). To assess whether EBF3 target genes are broadly dysregulated in APL, we examined their expression across orthogonal transcriptomic datasets. Gene expression changes of EBF3 targets in interferon and apoptosis pathways were broadly shared across *EBF3* knockdown, *PML::RARA* knockdown, and ATRA treatment in NB4 cells (Fig. 2J). The interferon gene targets but not apoptosis genes are lowly expressed in APL versus other AMLs in both BeatAML and TCGA LAML cohorts. These overlapping transcriptional data across cell line perturbations and patient samples reinforce EBF3 as a key effector of the PML::RARA transcriptional program.

### Recurrent mutations define a dosage corridor constraining EBF3

The essentiality of *EBF3* predicts that loss-of-function mutations would be deleterious in APL. Paradoxically, recurrent *EBF3* mutations occur in APL^24^ (13/348, 3.7%) but are absent in non-APL AML^22,23^ (0/789; Fisher’s exact p < 0.001; Fig. 3A). These mutations cluster in the DNA-binding domain and are predominantly truncating (10/13 nonsense or out-of-frame frameshift), predicting complete loss of function of the mutant allele (Fig. 3B). The remaining missense mutations (C198F, V315G) are predicted pathogenic by AlphaMissense^61^, suggesting impaired protein function rather than neutral variation.

**Figure 3.**
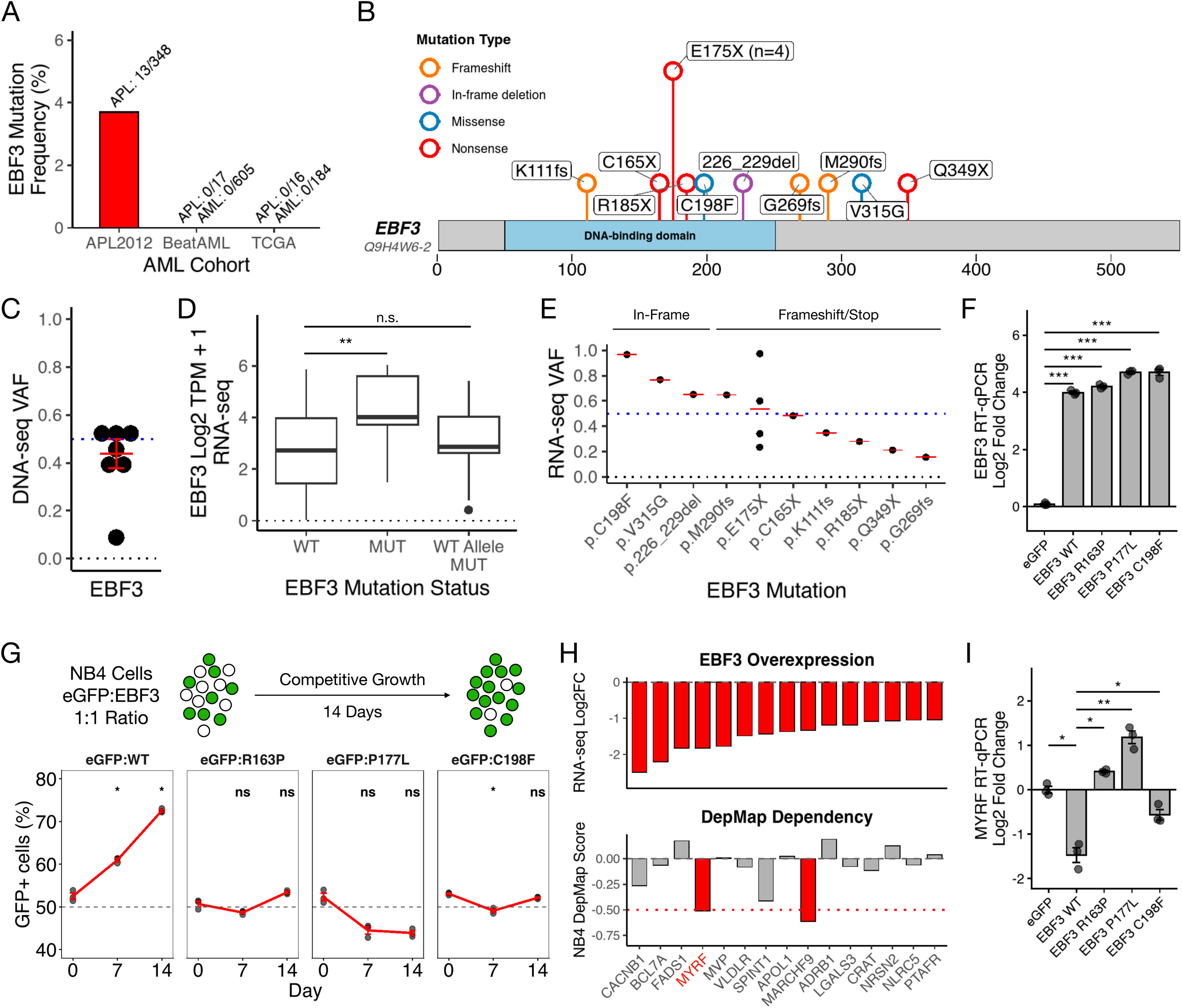
Recurrent *EBF3* mutations in APL mitigate growth suppression. (A) Frequency of *EBF3* mutations in APL versus non-APL AML across three patient cohorts (APL2012^24^, BeatAML^22^, TCGA-LAML^23^). (B) Lollipop diagram of the EBF3 protein (UniProt Q9H4W6-2) showing the positions of APL-derived mutations, which are largely clustered within the DNA-binding domain. X, stop codon. fs, frameshift. (C) Variant allele frequency (VAF) of the EBF3 mutations in the APL2012 cohort analyzed by whole exome sequencing (n = 7). (D) *EBF3* RNA expression (RNA-seq) of APL samples from the APL2012 cohort, quantified as Log2 TPM + 1. WT, APL with wild type EBF3 (n = 335). MUT, APL with mutant *EBF3* (n = 13). WT allele in MUT, wild type EBF3 allele expression in APL with mutant *EBF3* (n = 13). n.s., not significant. **, p < 0.01. Welch’s t-test. (E) VAFs of the *EBF3* mutations in the APL2012 cohort determined from RNA-seq, grouped by mutation type. (F) *EBF3* mRNA levels (RT-qPCR) of NB4 cells transduced with the annotated gene constructs. *ACTB* RNA was used as an internal control. n = 3. ***, p-value < 0.001. Welch’s t-test with Holm-Sidak adjustment. (G) Competitive proliferation assay. NB4 cells expressing *EBF3* wild type or mutants (C198F, R163P, P177L) were mixed 1:1 with eGFP-expressing control cells (identical backbone) and tracked over 14 days by flow cytometry. n = 3. ns, not significant. *, p < 0.05. Welch’s t-test with Holm-Sidak adjustment. (H) Top, EBF3 target genes with significant downregulation (FDR < 0.05, log2 fold change < -1) upon EBF3 overexpression as measured by RNA-seq. Below, NB4 DepMap scores of corresponding genes are shown below. MYRF dependency is unique to NB4 cells. (I) *MYRF* mRNA levels (RT-qPCR) of NB4 cells transduced with the annotated gene constructs. *ACTB* RNA was used as an internal control. n = 3. *, p-value < 0.05. **, p-value < 0.01. Welch’s t-test with Holm-Sidak adjustment.

If *EBF3* is essential, mutant cells must retain a functional wild-type allele. Indeed, all EBF3 mutations in the APL2012 cohort occur at variant allele frequencies (VAFs) around or below 0.5, consistent with preservation of the wild-type allele (Fig. 3C). Surprisingly, *EBF3*-mutant APL samples exhibit substantially higher total *EBF3* transcript levels than wild-type APL (Fig. 3D). RNA VAFs diverged from DNA VAFs in a mutation-type-dependent manner: truncating mutations generally showed reduced mutant allele expression consistent with nonsense-mediated decay, preserving wild-type transcript output (Fig. 3D-E). In contrast, in-frame mutations showed striking enrichment of the mutant allele at the RNA level (C198F DNA:RNA VAF 0.51:0.97; V315G DNA:RNA VAF 0.51:0.77). Because these in-frame mutations produce stable but functionally impaired protein, the elevated total *EBF3* expression in these samples largely reflects accumulation of hypomorphic transcript, while wild-type allele expression remains comparable to that observed in *EBF3* wild-type APL (Fig. 3D). This pattern suggests active selection, where cells tolerate reduced EBF3 function but cannot eliminate it entirely.

To investigate the functional significance of high mutant *EBF3* allele expression levels, we overexpressed wild-type EBF3 or EBF3 C198F in NB4 cells and measured the effect on cellular fitness compared to control NB4 cells overexpressing eGFP (Fig. 3F-G, Supplemental Fig. 3K). Forced overexpression of wild-type EBF3 reduced competitive fitness while EBF3 C198F did not. To understand why excess wild-type EBF3 decreases cell fitness, we performed RNA-seq on EBF3-overexpressing cells and intersected downregulated EBF3 target genes with NB4 essential genes from DepMap, nominating *MYRF* and *MARCHF9* as suppressed genes with phenotypic consequence (Fig. 3H). Critically, *MYRF* is a unique dependency for NB4 cells and not other myeloid cancer cell lines, and its expression is repressed upon wild-type EBF3 but not EBF3 C198F overexpression (Fig. 3I, Supplemental Fig. 3L). These findings indicate that while EBF3 is essential for APL cells, excessive EBF3 levels suppress growth by over-repressing genes required for APL survival and that *EBF3* mutations provide an escape from this growth suppression.

To confirm that loss of DNA-binding underlies escape from overexpression toxicity, we additionally tested HADDS-associated variants R163P and P177L, mutations within the DNA-binding domain previously shown to disrupt EBF3 DNA binding^29,30^ (Fig. 3F). Similar to C198F, both HADDS mutants did not suppress proliferation or repress *MYRF* (Fig. 3G, I). These data demonstrate that reduced EBF3 activity, whether through truncating mutations or hypomorphic alleles that impair EBF3 function, allows cells to escape the upper dosage constraint while retaining sufficient EBF3 wild type activity to support APL survival.

Together with the lethality of EBF3 loss (Fig. 2), these findings define a dosage corridor for EBF3 in APL: insufficient EBF3 (floor) triggers apoptosis via de-repression of cell death programs, while excess EBF3 (ceiling) represses essential genes required for proliferation. Heterozygous loss-of-function patient mutations represent an evolutionary solution to relax the ceiling without breaching the floor. This dosage corridor model, wherein both insufficient and excess EBF3 compromise fitness, provides a framework for understanding EBF3’s constrained role in APL. Critically, the mutations that result carry a tumor suppressor signature yet mark an essential gene – a pattern that would be misinterpreted by standard classification.

### PML::RARA enforces EBF3 expression via a p300-dependent enhancer

As EBF3 expression is associated with PML::RARA status and PML::RARA is a chromatin-binding factor, we examined the chromatin regulatory landscape at the *EBF3* gene locus. PML::RARA binding was detected within an intron of the *EBF3* gene but not at the promoter/transcription start site (TSS) region (Fig. 4A). At both regions, we observed RNA polymerase II (RNAPII) occupancy and accessible chromatin, shared in both NB4 and primary APL cells (Fig. 4A). ATRA treatment dramatically reduced RNAPII occupancy across the *EBF3* gene, whereas accessible chromatin did not change as dramatically. Intriguingly, in U937-PR9 cells harboring a zinc-inducible *PML::RARA* transgene^14,62^, the fusion binds the *EBF3* intronic enhancer upon induction, yet *EBF3* transcripts are not detected (Supplemental Fig. 4A-B). This likely reflects the absence of accessible chromatin at the *EBF3* TSS in this cellular context, suggesting that PML::RARA binding alone is insufficient without a permissive chromatin state.

**Figure 4.**
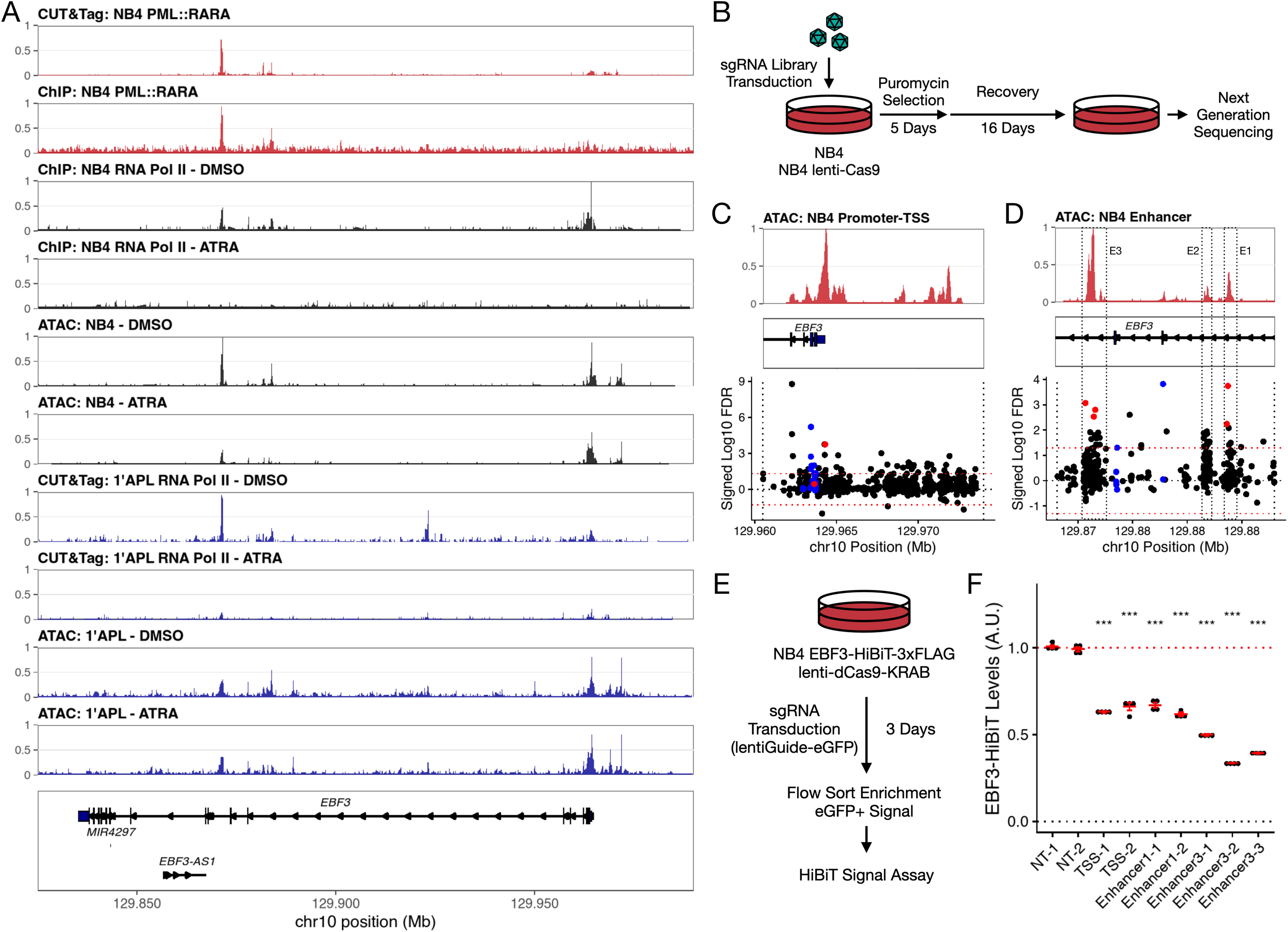
PML::RARA activates EBF3 via an essential intronic enhancer. (A) Genome browser view of PML::RARA binding^17,18^ at the *EBF3* gene locus in NB4 cells. RNA polymerase II binding and chromatin accessibility upon 48 hours of 1 μM ATRA or DMSO treatment in NB4 cells^45,76^ or a primary APL patient sample are also shown. (B) Schematic of the CRISPR/Cas9 *EBF3* gene tiling viability screen. (C) CRISPR/Cas9 sgRNA enrichment at the *EBF3* promoter/transcription start site (TSS). Higher y-values represent increased sgRNA dropout. Chromatin accessibility profile is shown for reference. Blue, sgRNAs targeting *EBF3* protein-coding sequence. Red, tested sgRNAs in Figure 4E-F. (D) CRISPR/Cas9 sgRNA enrichment at the *EBF3* intronic enhancer. Higher y-values represent increased sgRNA dropout. Chromatin accessibility profile is shown for reference. E1-3, enhancers 1-3. Blue, sgRNAs targeting *EBF3* protein-coding sequence. Red, tested sgRNAs in Figure 4E-F. (E) Schematic of CRISPR interference-based validation of *EBF3* gene regulatory elements. (F) EBF3-HiBiT levels (NanoBiT lytic assay) of eGFP+ cells sorted 3 days post-transduction. NT, nontargeting sgRNA. All statistical comparisons were made relative to NT-2. n = 4. ***, p < 0.001. Welch’s t-test with Holm-Sidak adjustment.

To functionally map essential regulatory elements, we performed a CRISPR/Cas9 tiling screen across the *EBF3* locus^63^. Because disruption of regulatory elements would reduce *EBF3* expression below the viability threshold, we used sgRNA depletion as a proxy for functional importance (Fig. 4B, Supplemental Fig. 4C-E). In addition to sgRNAs targeting protein-coding sequence, sgRNAs targeting both promoter/TSS and the intronic enhancer were depleted (Fig. 4C-D, Supplemental Fig. 4F). Importantly, dCas9-KRAB (CRISPR interference^64^)-mediated repression of either the intronic enhancer or the TSS reduced *EBF3* levels, confirming their functional requirement for gene regulation (Fig. 4E-F). Altogether, these data indicate that PML::RARA binds at an intronic enhancer within the *EBF3* gene to activate *EBF3* gene expression.

The dependence of EBF3 expression on an active enhancer suggested a pharmacologic strategy: inhibiting enhancer activity to suppress *EBF3*. p300 (EP300) is a histone acetyltransferase required for enhancer function and is itself an essential gene in NB4 cells. p300 is present at the PML::RARA-bound *EBF3* intronic enhancer alongside H3K27Ac, its enzymatic target, in NB4 cells^18^; likewise, patient APL exhibit high H3K27Ac signal at this region^65^ (Fig. 5A, Supplemental Fig. 5A-B). Treatment with A485^66^, a potent and selective p300 inhibitor, leads to decreased viability, EBF3 protein and RNA levels, and H3K27Ac levels in a dose-dependent manner, with 72-hour IC50 around 120 nM (Supplemental Fig. 5C-E). A485 reduced EBF3 protein levels prior to any decrease in cell viability, consistent with EBF3 loss as a cause rather than consequence of cell death (Fig. 5B-C, Supplemental Fig. 5F). Similar sub-micromolar IC50 (185 nM) values and decreased *EBF3* RNA and H3K27Ac levels were observed in A485-treated NB4-R2 cells (Supplemental Fig. 5C, G-H). To test whether *EBF3* suppression contributes to A485-mediated cell death, we treated NB4 cells expressing exogenous *EBF3* with A485. Indeed, EBF3 but not eGFP overexpression rescued viability at 0.03 and 0.1 µM A485 after 6 days of treatment, concentrations at which exogenous EBF3 expression was maintained (Fig. 5D, Supplemental Fig. 5I-J). These data demonstrate that the EBF3 dosage corridor creates a pharmacologically actionable vulnerability: p300 inhibition pushes EBF3 below the viability threshold, exploiting the same constraint that patient mutations navigate from above.

**Figure 5.**
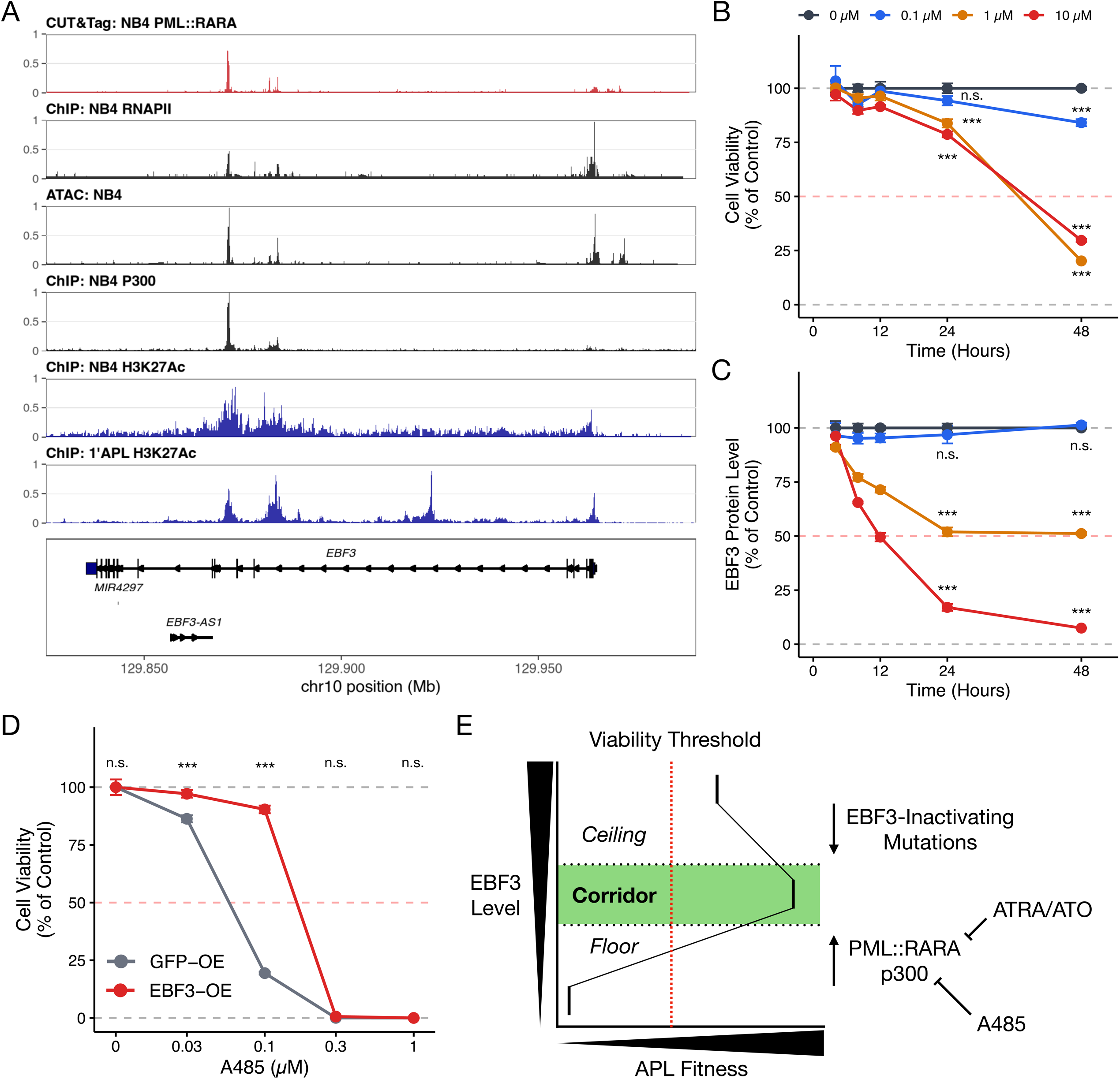
EBF3 dependency is therapeutically exploitable via p300 inhibition. (A) Genome browser view of p300 and H3K27ac enrichment in NB4 cells^18^ and mean enrichment of H3K27Ac in primary APL patient samples^65^ are also shown. PML::RARA, RNA polymerase II, and chromatin accessibility enrichment in NB4 are shown for reference. (B) CellTiter-Glo viability of NB4 cells treated with varying concentrations of A485 for the indicated time. Luminescence is normalized to the corresponding 0 μM condition. n = 3. n.s., not significant. ***, p < 0.001. Dunnett’s test (vs. 0 μM control). (C) EBF3-HiBiT luminescence (NanoBiT lytic assay) normalized by corresponding CellTiter-Glo viability of NB4 cells treated with varying concentrations of A485 for the indicated time. Luminescence ratio is normalized to the corresponding 0 μM condition. n = 3. n.s., not significant. ***, p < 0.001. Dunnett’s test (vs. 0 μM control). (D) CellTiter-Glo viability of NB4 cells harboring constitutive *eGFP-FLAG* (*GFP-OE*) or FLAG-*EBF3* (*EBF3-OE*) transgenes expressed via an EF1alpha promoter and treated for 6 days at the indicated concentrations of A485. Luminescence is normalized to the corresponding 0 μM condition. n = 4. n.s., not significant. ***, p < 0.001. Pairwise comparison with Sidak correction. (E) Schematic model of EBF3 as a corridor TF. PML::RARA and p300 enforce EBF3 expression above the floor to maintain the APL cell state, which can be suppressed by ATRA/ATO targeting PML::RARA or A485 inhibiting p300. High EBF3 is selected against via the emergence of heterozygous *EBF3*-inactivating mutations, thereby optimizing APL cell proliferation.

## Discussion

Our findings demonstrate that oncogenic fusions can impose dosage constraints that convert genes with tumor suppressor activity into essential dependencies, a relationship that subverts conventional mutation-based classification. The resulting mutations carry a tumor suppressor signature but indicate the opposite: essential genes under adaptive constraint. We show that PML::RARA creates such a constraint on EBF3 in APL (Fig. 5E). PML::RARA activates EBF3 through a p300-dependent intronic enhancer, creating a dependency that is genetically essential, evolutionarily constrained, and pharmacologically actionable.

The transcriptional correlation between *EBF3* knockdown and *PML::RARA* knockdown suggests that EBF3 is not merely one of many PML::RARA targets, but a major effector of the fusion’s transcriptional program. PML::RARA exerts oncogenic activity through downstream transcription factor networks, both by suppressing differentiation-promoting factors (e.g., PU.1/SPI1, CEBPA)^38,39^ and by restraining cell death programs^67,68^. EBF3 appears to mediate the latter: direct EBF3 targets include apoptosis and interferon response genes that are de-repressed upon EBF3 loss. Interferon signaling functions as a tumor suppressive mechanism in APL, inducing leukemic stem cell exhaustion and apoptosis^69,70^. EBF3-mediated repression of these programs may thus be essential for leukemia-initiating cell survival. Other transcription factors such as KLF4 and GATA2 also contribute to APL transformation^71,72^, suggesting EBF3 operates within a broader network of PML::RARA-regulated factors.

Why does PML::RARA drive EBF3 to high levels despite the constraints this creates? We propose that EBF3-mediated repression of apoptosis and interferon programs is necessary for APL survival, requiring PML::RARA to activate EBF3 above a minimum threshold. Additionally, interferon signaling can reshape the bone marrow microenvironment, and EBF3-mediated suppression of these programs may further support a niche permissive for leukemic maintenance. However, the same transcriptional activity that represses death genes also silences essential genes such as *MYRF* when EBF3 levels rise too high. The corridor thus reflects a fundamental tradeoff: sufficient EBF3 to repress death programs, but not so much as to silence essential genes. Consistent with this model, PML::RARA appears to calibrate EBF3 expression near the upper boundary of this corridor, creating selective pressure for mutations that reduce activity toward the corridor’s center.

This dosage-constrained dependency distinguishes *EBF3* from conventional oncogenes and tumor suppressors^3^. In solid tumors and pediatric AML, EBF3 overexpression induces apoptosis and cell cycle arrest, consistent with tumor suppressor activity^32,33^. In APL, our data reveal a distinct mechanism: excess EBF3 over-represses essential genes such as *MYRF* rather than activating death programs directly. The resulting fitness cost explains why heterozygous loss-of-function mutations cluster in the EBF3 DNA-binding domain in APL but not in other AMLs^22–24^. By comparing an APL-derived mutation (C198F) with germline HADDS mutations (R163P, P177L)^30^, we demonstrate that all three attenuate EBF3 transcriptional activity, allowing cells to escape the growth-suppressive ceiling while retaining sufficient function to survive.

The mechanism by which PML::RARA activates EBF3 via an intronic enhancer is consistent with recent work demonstrating that PML::RARA acts as a transcriptional activator^17,18,73^. PML::RARA recruits p300 to establish super-enhancers^74,75^ at genes including *GFI1*, a key gene required for leukemic maintenance. Our findings extend this framework: PML::RARA binding at the *EBF3* intronic enhancer, together with p300 recruitment and H3K27 acetylation, drives EBF3 expression in APL. However, this mechanism appears to require a permissive chromatin state at the TSS: in U937-PR9 cells, PML::RARA binds the *EBF3* intronic enhancer yet fails to activate transcription, coincident with closed chromatin at the promoter. This may explain why EBF3 was not identified as a PML::RARA target in prior studies using this model system^17,76,77^.

The dependence of EBF3 on p300-mediated enhancer activity creates a therapeutic opportunity^66^: A485 suppresses EBF3 and kills APL cells, including ATRA-resistant NB4-R2, with EBF3 overexpression providing partial rescue. Conservation of EBF3 expression across resistant APL and rare RARA fusion subtypes (e.g., PLZF::RARA^52^, FIP1L1::RARA^53,54^, STRN3::RARA^55^) suggests this vulnerability may extend beyond treatment-sensitive PML::RARA-driven disease^56,78^.

More broadly, the dosage corridor, wherein essential genes are constrained from above as well as below, may apply to other regulatory factors in fusion-driven cancers. A dosage corridor gene is defined by three features: vulnerability upon knockdown, aberrant overexpression imposed by oncogenic context, and fitness loss upon overexpression. Genes with pleiotropic, dose-dependent effects on cell state, such as transcription factors, kinases, or microRNAs, may be particularly susceptible to such constraints^79^. The heterozygous loss-of-function mutations we observe in EBF3 are a consequence of this corridor. Indeed, this mutation pattern (heterozygous, clustered in functional domains, and present only in specific oncogenic contexts) may serve as a signature to nominate candidate corridor genes in other cancers. Conservation of EBF3 expression across RARA fusion-driven APL subtypes suggests the corridor mechanism may be a general feature of RARA fusion oncoproteins. Whether non-fusion oncogenes impose analogous constraints remains to be determined, though any oncogene that aberrantly activates a pleiotropic transcription factor could in principle create similar dosage tradeoffs.

Dosage corridor genes share features with developmental “Goldilocks” factors but arise from a distinct mechanism^80,81^. Intrinsic dosage sensitivity reflects a gene’s inherent properties, whereas dosage corridors are imposed by oncogenic context. In APL, PML::RARA pushes EBF3 toward a ceiling constraint. The mutations we observe are not evidence of intrinsic haploinsufficiency but rather adaptations to an oncogene-created problem.

These findings carry implications for cancer genomics^82^: heterozygous loss-of-function mutations are conventionally interpreted as evidence of tumor suppression^4^, yet corridor mutations indicate the opposite. Tumor suppressor-like mutations marking essential genes are signatures of adaptive change that emerge within oncogene-imposed constraints. Systematic identification of fusion-imposed dosage constraints may reveal dependencies currently misclassified as tumor suppressors.

## Acknowledgements

We are grateful to the Harvard Medical School Biopolymers Facility, Harvard Medical School Research Computing, and the BIDMC flow cytometry core for their assistance and support. We also thank Dr. Steven P. Gygi and the Taplin Mass Spectrometry Facility at Harvard Medical School for use of their mass spectrometers and access to their COMET based data analysis suite. We thank Arne Müller, Dr. Cristina Bogsan, Dr. Tanvi Saxena, Dr. Christos Miliotis, Dr. Yu-Ru Lee, Dr. Priscilla Cheung, and all members of the Slack laboratory for their input and support. We thank all funding sources listed below for their support. We especially thank the Chiles family, the Hope Funds for Cancer Research Foundation, and the Wipe Out Kids’ Cancer Foundation for their support to J.D.L.

## Funding Sources

Hope Funds for Cancer Research (J.D.L.)

Wipe Out Kids’ Cancer Foundation (J.D.L.)

NIH/NCI K99 CA293268 (J.D.L.)

Harvard Ludwig Cancer Center (F.J.S.)

NIH/NCI R35 CA232105 (F.J.S.)

NIH/NIGMS R01 GM132129 (J.A.P.)

Harvard College Research Program (F.L., E.H.S.)

FIS 3 FIS-2024-06156 project (M.T.V.)

NIH/NHLBI P01 HL131477 (D.G.T.)

## Author Contributions

Conceptualization: J.D.L., M.M.U., F.J.S.

Investigation: J.D.L., M.M.U., F.L., R.R.P., E.H.S., A.C.H., T.O., G.C., D.B.C. Analysis: J.D.L., R.R.P., J.A.P.

Resources: J.D.L., D.B.C., B.Q.T., J.G.C., D.G.T., M.T.V., T.A.G., F.J.S.

Supervision: J.D.L., F.J.S.

Writing: original draft: J.D.L., M.M.U., F.J.S.

## Supplemental Figure Legends

**Supplemental Figure 1.**
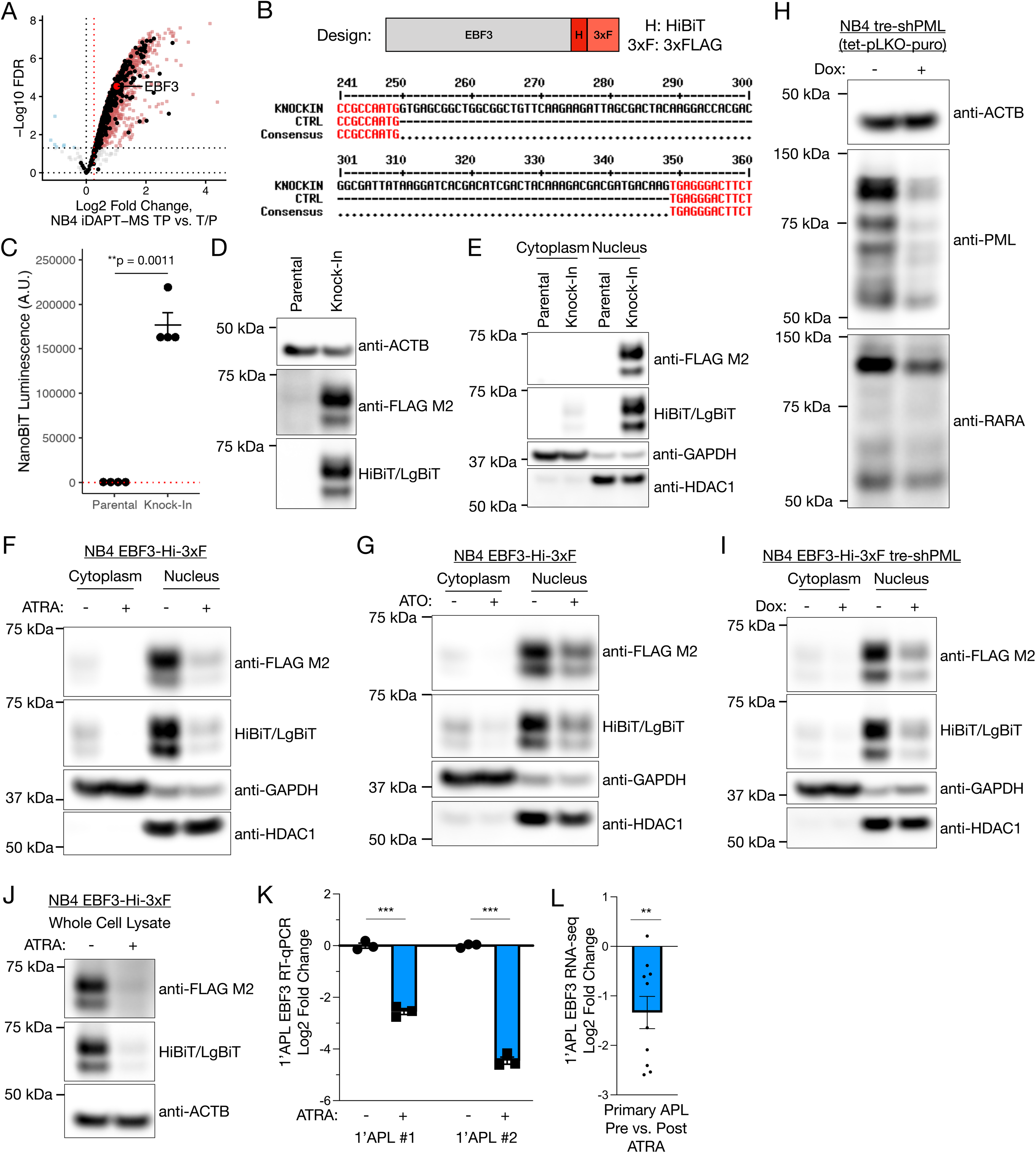
Extended characterization of EBF3 as an APL-specific factor. (A) iDAPT-MS profiling of NB4 cells^45^ tagged with transposase-peroxidase fusion (TP; n = 3) versus transposase only or peroxidase only (T/P; n = 4), representing background signal. Black points, CisBP transcription factors^85^. (B) Top, design of the CRISPR knock-in tagging strategy. Bottom, Sanger sequencing validation of *EBF3* knock-in in NB4 cells. Inference of CRISPR Edits (ICE) analysis shows 100% knock-in efficiency. (C) EBF3 protein expression (NanoBiT lytic assay) in parental or knock-in NB4 cells. n = 4. **, p < 0.01. Welch’s t-test. (D) EBF3 protein expression (Western blotting) in parental or knock-in NB4 cells. (E) EBF3 nuclear and cytoplasmic expression (Western blotting) in parental or knock-in NB4 cells. (F) EBF3 nuclear and cytoplasmic expression (Western blotting) in EBF3 knock-in NB4 cells following 1 μM ATRA or vehicle treatment for 48 hours. (G) EBF3 nuclear and cytoplasmic expression (Western blotting) in EBF3 knock-in NB4 cells following 1 μM arsenic trioxide (ATO) or vehicle treatment for 48 hours. (H) PML and RARA protein expression (Western blotting) in NB4 cells following inducible PML::RARA shRNA knockdown with or without 1 μg/mL doxycycline for 72 hours. (I) EBF3 nuclear and cytoplasmic expression (Western blotting) in EBF3 knock-in NB4 cells following PML::RARA shRNA knockdown with or without 1 μg/mL doxycycline for 72 hours. (J) EBF3 protein expression (Western blotting) in knock-in NB4 cells following 1 μM ATRA or vehicle treatment for 48 hours by Western blotting. (K) *EBF3* mRNA expression (RT-qPCR) in primary APL cells cultured *ex vivo* following 1 μM ATRA or vehicle treatment for 48 hours. n = 3. ***, p < 0.001. Welch’s t-test. (L) *EBF3* RNA expression (RNA-seq) of primary APL samples harvested before and after 48 hours of ATRA treatment in patients (*in vivo*)^25^. n = 10. **, p < 0.01. One-sample t-test.

**Supplemental Figure 2.**
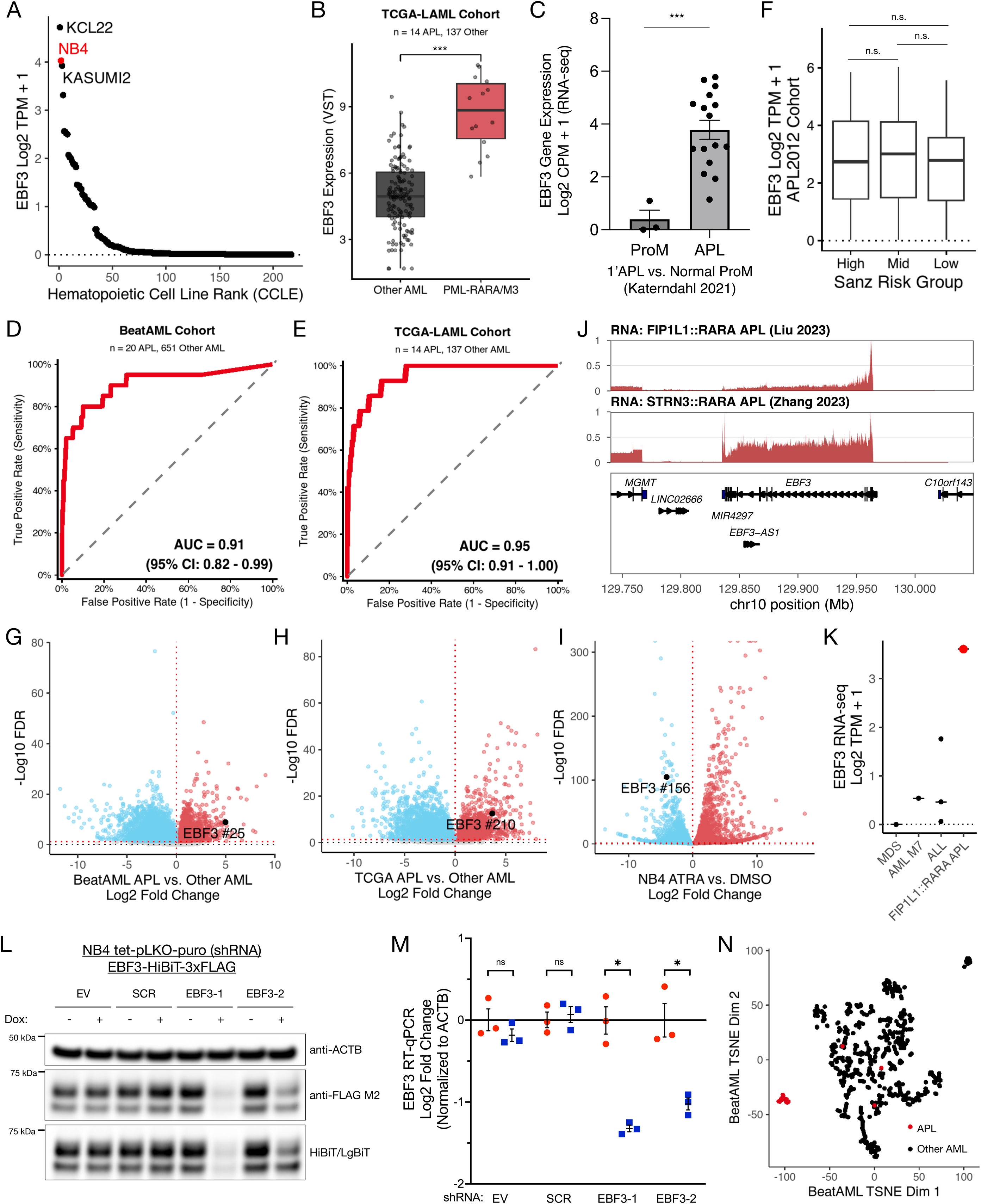
Extended characterization of EBF3 expression and dependency. (A) *EBF3* RNA expression (RNA-seq) of hematopoietic cancer cell lines in the Cancer Cell Line Encyclopedia^47,48^, quantified as Log2 TPM + 1. n = 217. (B) *EBF3* RNA expression (RNA-seq) in APL patient samples compared to other AML subtypes in the TCGA-LAML cohort^23^, quantified as VST (variance stabilizing transform) values. ***, p < 0.001. Wilcoxon rank-sum test. (C) *EBF3* RNA expression (RNA-seq) in APL patient samples (n = 16) compared to normal promyelocytes^50^ (ProM; n=3), quantified as log2 CPM + 1. ***, p < 0.001. Welch’s t-test. (D) Receiver operating characteristic (ROC) curve for *EBF3* expression as a classifier of APL versus other AML subtypes (BeatAML cohort). (E) Receiver operating characteristic (ROC) curve for *EBF3* expression as a classifier of APL versus other AML subtypes (TCGA-LAML cohort). (F) *EBF3* RNA expression (RNA-seq) of APL samples stratified by Sanz clinical risk group (High: n = 104; Mid: n = 157; Low: n = 62) from the APL2012 cohort^24^, quantified as Log2 TPM + 1. n.s., not significant. Welch’s t-test. (G) Volcano plot of gene expression differences between APL and other AML in the BeatAML cohort. (H) Volcano plot of gene expression differences between APL and other AML in the TCGA-LAML cohort. (I) Volcano plot of gene expression changes of NB4 cells upon ATRA versus DMSO treatment. (J) Genome track view of RNA expression (RNA-seq) at the *EBF3* gene locus of primary APL driven by rare RARA fusions (FIP1L1::RARA t(4;17)^54^, STRN3::RARA t(14;17)^55^). (K) *EBF3* RNA expression (RNA-seq) of leukemia samples from the Liu 2023 dataset^54^, quantified as Log2 TPM + 1. (L) FLAG and HiBiT (EBF3) protein expression (Western Blotting) of NB4 knock-in cells harboring doxycycline-inducible *EBF3* or negative control shRNA transgenes (SCR, scrambled; EV, empty vector) and treated with or without 1 μg/mL doxycycline for 72 hours. (M) *EBF3* mRNA expression (RT-qPCR) of NB4 cells harboring doxycycline-inducible *EBF3* or negative control shRNA transgenes (SCR, scrambled; EV, empty vector) and treated with or without 1 μg/mL doxycycline for 72 hours. *ACTB* RNA was used as an internal control. n = 3. n.s., not significant. *, p < 0.05. Welch’s t-test. (N) t-SNE projection of BeatAML RNA-seq profiles using significant differentially expressed genes (FDR < 0.05, |Log2 Fold Change| > 1) upon shEBF3 knockdown as features.

**Supplemental Figure 3.**
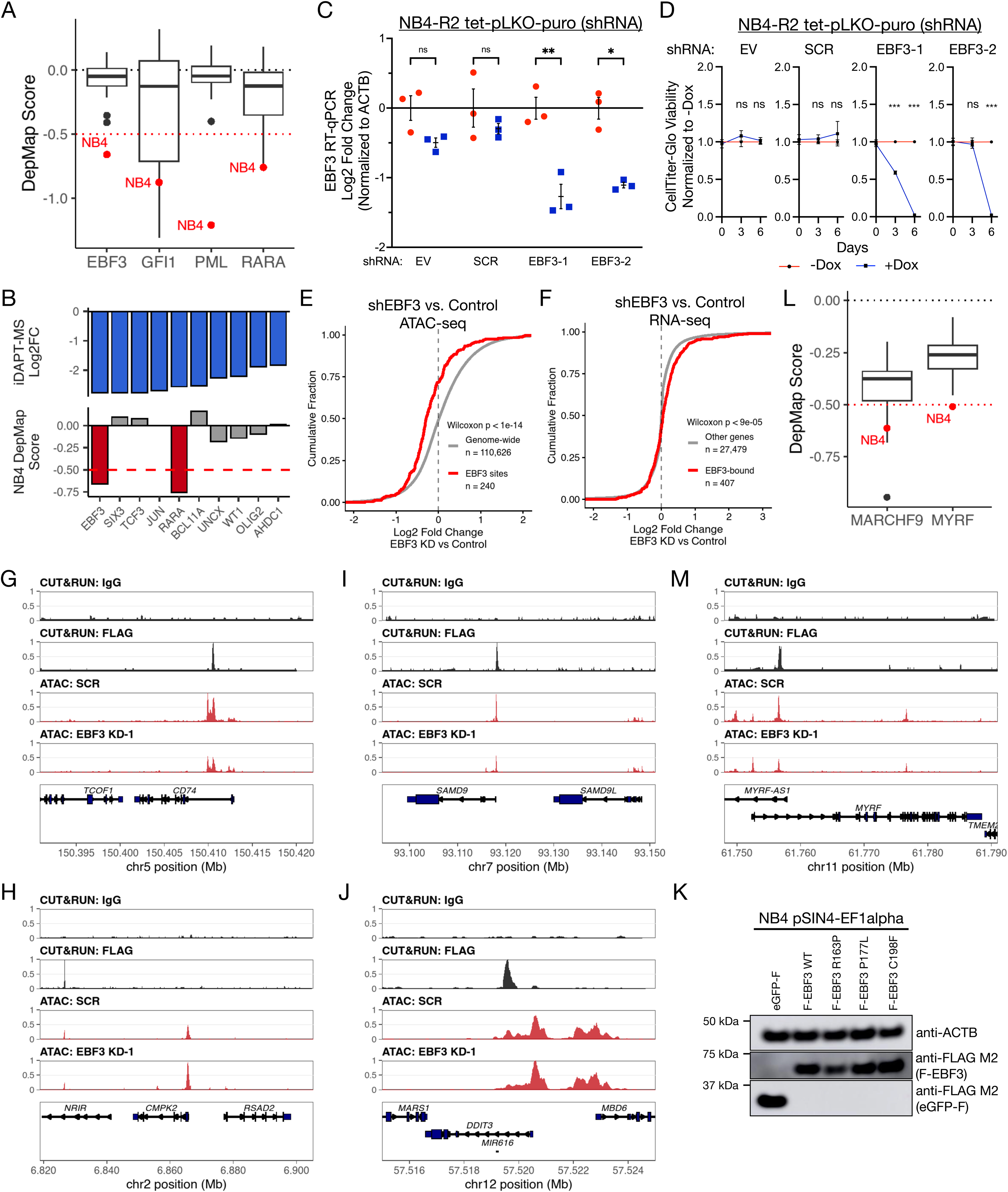
Extended characterization of EBF3 transcriptional and chromatin regulation. (A) DepMap 23Q4^57,58^ Chronos dependency scores of EBF3, GFI1, PML, and RARA across myeloid cancer cell lines (CRISPR/Cas9 knockout-based). EBF3, PML, and RARA dependencies are strongest in NB4 cells relative to other myeloid cancer cell lines. (B) Top, top 10 iDAPT-MS proteins lost upon ATRA treatment as in Fig. 1B. Bottom, DepMap dependency scores corresponding to these proteins in NB4 cells. (C) *EBF3* mRNA expression (RT-qPCR) of NB4-R2 cells harboring doxycycline-inducible EBF3 or negative control shRNA transgenes (SCR, scrambled; EV, empty vector) and treated with or without 1 μg/mL doxycycline for 72 hours. *ACTB* RNA was used as an internal control. n = 3. n.s., not significant. *, p < 0.05. **, p < 0.01. Welch’s t-test. (D) CellTiter-Glo viability of NB4-R2 cells harboring doxycycline-inducible EBF3 or negative control shRNA transgenes (SCR, scrambled; EV, empty vector) and treated with or without 1 μg/mL doxycycline for the indicated time. Luminescence is normalized to the corresponding -Dox condition. n = 5. ns, not significant. ***, p < 0.001. Welch’s t-test. (E) Distribution of chromatin accessibility changes upon *EBF3* knockdown genome-wide or at EBF3-binding sites. (F) Distribution of gene expression changes upon *EBF3* knockdown transcriptome-wide or at genes proximal to EBF3 binding (within 50 kb). (G) Genome browser view of the CD74 locus showing EBF3 (FLAG) binding and *EBF3* knockdown chromatin accessibility changes. (H) Genome browser view of the CMPK2/RSAD2 locus showing EBF3 (FLAG) binding and *EBF3* knockdown chromatin accessibility changes. (I) Genome browser view of the SAMD9/SAMD9L locus showing EBF3 (FLAG) binding and *EBF3* knockdown chromatin accessibility changes. (J) Genome browser view of the *DDIT3* locus showing EBF3 (FLAG) binding and *EBF3* knockdown chromatin accessibility changes. (K) FLAG protein expression (Western blotting) of NB4 cells with constitutive FLAG-EBF3 variants or eGFP-FLAG. (L) DepMap 23Q4 Chronos dependency scores of *MARCHF9* and *MYRF* across myeloid cancer cell lines (CRISPR/Cas9 knockout-based). MYRF dependency is strongest in NB4 cells relative to other myeloid cancer cell lines. (M) Genome browser view of the *MYRF* locus showing EBF3 (FLAG) binding and EBF3 knockdown chromatin accessibility changes.

**Supplemental Figure 4.**
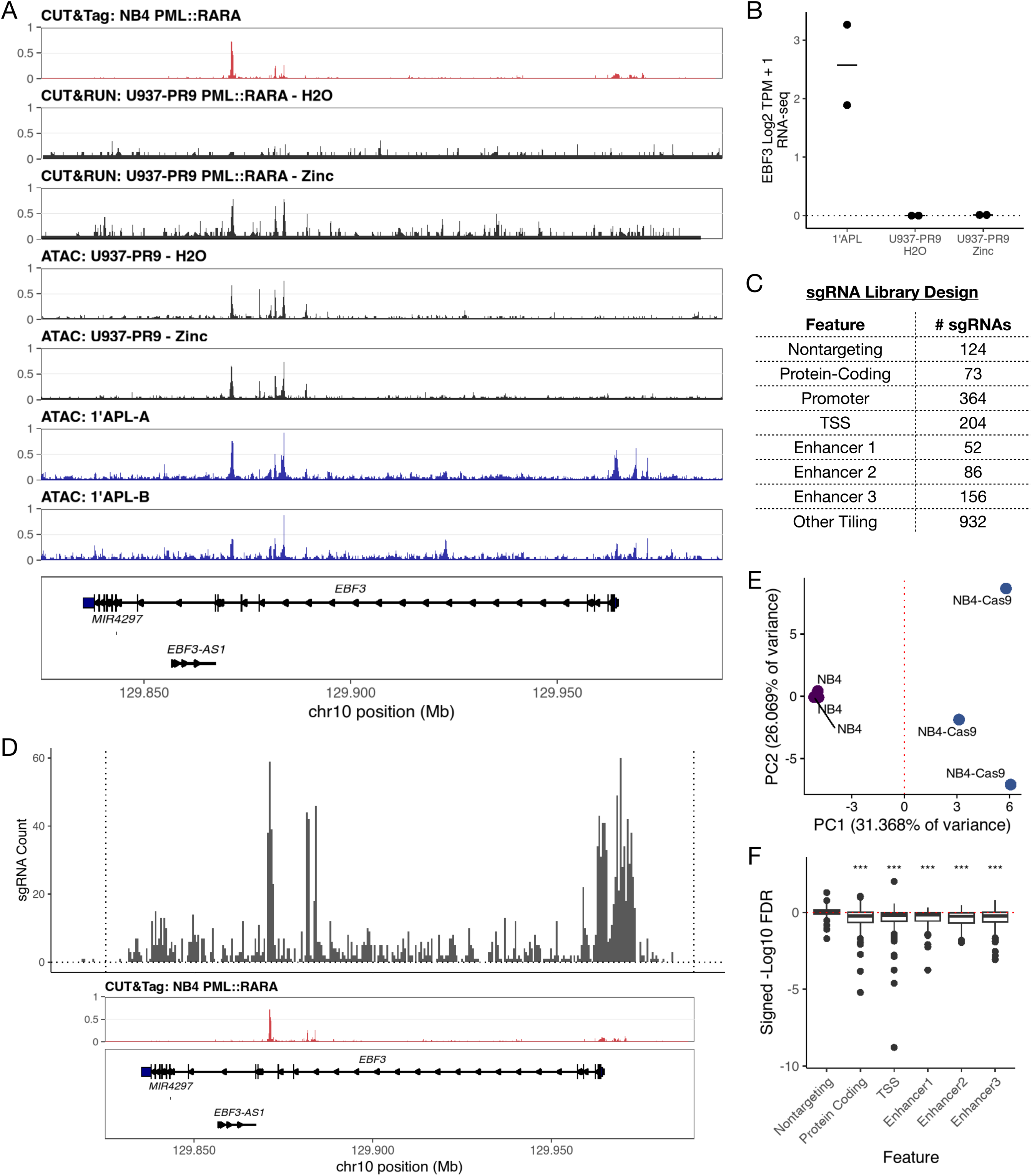
Extended characterization of EBF3 locus regulation. (A) Genome browser view of the *EBF3* locus showing PML::RARA and chromatin accessibility enrichment with or without zinc in U937-PR9 cells and chromatin accessibility enrichment of two patient APL samples from Villiers 2023^17^. PML::RARA enrichment in NB4 is shown for reference. (B) *EBF3* RNA expression (RNA-seq) of U937-PR9 cells treated with or without zinc and two patient APL samples from Villiers 2023^17^, quantified as Log2 TPM + 1. (C) sgRNA library design and targeted *EBF3* genic features. (D) Distribution of CRISPR/Cas9 tiling sgRNAs across the *EBF3* gene locus. PML::RARA enrichment in NB4 is shown below for reference. (E) Principal component analysis of sgRNA distribution counts with or with Cas9 expression. (F) Feature-specific sgRNA enrichment from the CRISPR/Cas9 EBF3 tiling screen (TSS [n = 204], enhancer [E1, n = 52; E2, n = 86; E3, n = 156], protein-coding [n = 73], nontargeting controls [n = 124]). CRISPR/Cas9 targeting of the PML::RARA-bound intronic enhancer or the promoter/TSS cause loss of cell viability. ***, p < 0.001. Welch’s t-test with Holm-Sidak adjustment.

**Supplemental Figure 5.**
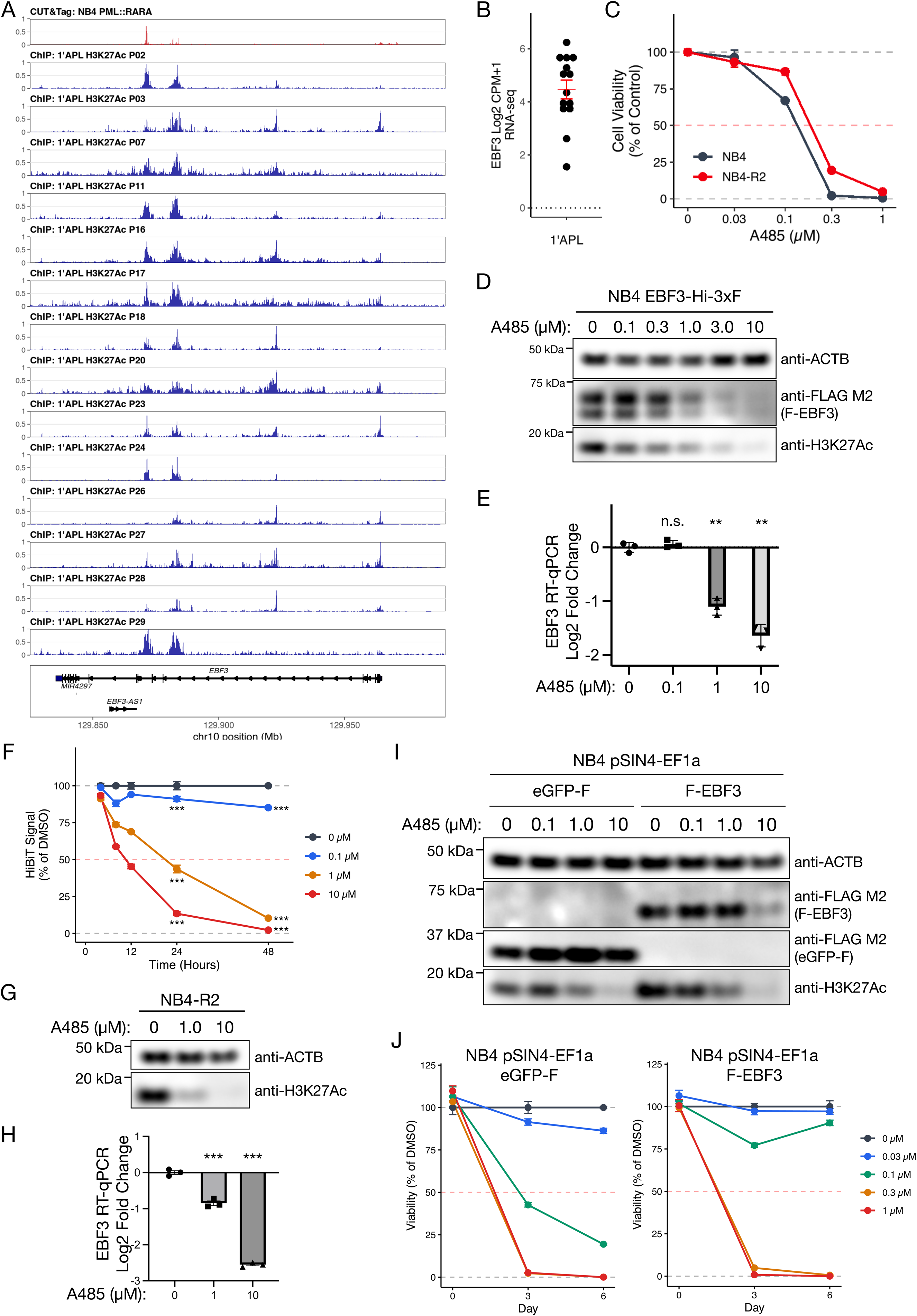
Extended characterization of A485 response in APL cells. (A) Genome browser view of H3K27Ac ChIP-seq enrichment of individual patient APL samples^65^, corresponding to Figure 5A. PML::RARA enrichment in NB4 cells is shown for reference. (B) *EBF3* RNA expression (RNA-seq) of primary APL samples quantified as Log2 CPM + 1^65^, corresponding to Supplemental Figure 5A. (C) CellTiter-Glo viability of NB4 and NB4-R2 cells treated with varying doses of A485 for 72 hours. n = 4. NB4: IC50 = 120 nM (95% CI: 101-139 nM). NB4-R2: IC50 = 185 nM (95% CI: 161-209) (D) EBF3 and H3K27Ac protein expression (Western blotting) of knock-in NB4 cells treated with varying doses of A485 for 24 hours. (E) *EBF3* RNA expression (RT-qPCR) of NB4 cells treated with varying doses of A485 for 24 hours. *ACTB* RNA was used as an internal control. n = 3. n.s., not significant. **, p < 0.01. Welch’s t-test. (F) EBF3-HiBiT luminescence (NanoBiT lytic assay) of NB4 cells treated with varying concentrations of A485 for the indicated time. Luminescence ratio is normalized to the corresponding 0 μM condition. n = 3. ***, p < 0.001. Dunnett’s test (vs. 0 μM control). (G) H3K27Ac protein expression (Western blotting) of NB4-R2 cells treated with varying doses of A485 for 24 hours. (H) *EBF3* RNA expression (RT-qPCR) of NB4-R2 cells treated with varying doses of A485 for 24 hours. *ACTB* RNA was used as an internal control. n = 3. ***, p < 0.001. Welch’s t-test. (I) FLAG and H3K27Ac protein expression (Western blotting) of NB4 cells with constitutive FLAG-EBF3 or eGFP-FLAG expression following varying doses of A485 after 24 hours. (J) CellTiter-Glo viability of NB4 cells harboring constitutive *eGFP-FLAG* (*GFP-OE*) or FLAG- *EBF3* (*EBF3-OE*) transgenes following varying doses of A485 over 6 days. Luminescence is normalized to the corresponding 0 μM condition. n = 4. GFP-OE vs. EBF3-OE at 0.1 μM: ***p < 0.001 at Day 3 and Day 6. Pairwise comparison with Sidak correction.

## Supplementary Tables

Table S1. Dataset summary. (A) Post-processed publicly available datasets. (B) Publicly available raw sequencing datasets.

Table S2. Differential gene expression analyses. (A) NB4 shEBF3 versus control. (B) NB4 shPML versus control. (C) NB4 EBF3 versus eGFP overexpression.

Table S3. EBF3 CUT&RUN peaks and annotations

Table S4. Pathway enrichment analyses. (A) NB4 shEBF3 versus control. (B) NB4 shPML versus control.

Table S5. APL2012 cohort EBF3 mutation information and corresponding datasets. Table S6. CRISPR tiling screen analyses.

Table S7. Oligonucleotide and primer sequences.

## Materials and Methods

### Resources and Models

#### Cell Lines and Culture Conditions

NB4 cells were obtained from DSMZ (ACC 207). NB4-R2 cells were obtained from the Tenen lab. Both cell lines were maintained in RPMI 1640 medium (Gibco) supplemented with 10% fetal bovine serum (FBS; Sigma) and 1% penicillin/streptomycin (Thermo Fisher Scientific). For experiments involving tetracycline/doxycycline-inducible constructs, cells were cultured with 10% tetracycline-free FBS (Takara). HEK293T cells (ATCC, CRL-3216) were maintained in DMEM (Gibco) supplemented with 10% FBS and 1% penicillin/streptomycin. All cells were grown at 37°C in a humidified incubator with 5% CO₂.

#### Primary Patient Samples

*RT-qPCR validation cohort.* Primary bone marrow samples from leukemia patients were obtained at Tor Vergata University with informed patient consent approval by the Tor Vergata Ethical Committee (Study Protocol NM-MY STUDY – Clinical Trial Registry 34/20). Total RNA was extracted from mononuclear cells (MNCs), isolated by Ficoll-Hypaque density-gradient centrifugation, using TRIzol Reagent (Invitrogen, Thermo Fisher Scientific, Waltham, MA, USA) according to the manufacturer’s recommendations, and reverse transcribed into cDNA as previously described (PMID: 37803061). The resulting cDNAs were used for RT-qPCR validation described below.

*Ex vivo functional studies.* Peripheral blood or bone marrow aspirates were obtained from APL patients at Massachusetts General Hospital with written informed consent under a protocol approved by the Institutional Review Board at the Dana-Farber/Harvard Cancer Center (Protocol #13-583). Samples were subjected to red cell lysis, and the buffy coat was isolated and cryopreserved in 10% DMSO with autologous serum. For ex vivo studies, cells were thawed and cultured for up to 7 days in IMDM (Gibco) supplemented with 20% FBS, 1% penicillin/streptomycin, and the following cytokines (BioLegend): SCF (50 ng/mL), G-CSF (50 ng/mL), IL-6 (20 ng/mL), and IL-3 (10 ng/mL). After 24-48 hours, dead cells were removed using the Dead Cell Removal Kit (Miltenyi Biotec) according to the manufacturer’s instructions, and the resulting live cells were used for subsequent experiments.

#### Mouse Studies

All animal experiments were approved by the Institutional Animal Care and Use Committee at Beth Israel Deaconess Medical Center and conducted in accordance with institutional guidelines. Male Balb/c nude mice (Jackson Laboratory) were used for xenograft studies and housed under specific pathogen-free conditions. NB4 cells (5 × 10⁶) transduced with doxycycline-inducible shRNA constructs were injected subcutaneously into the flank (n = 10 per group). Tumor sizes were monitored by caliper measurement. After 15 days, when tumors were palpable, mice were placed on doxycycline-containing chow (2.3 g/kg; Bio-Serv) to induce shRNA expression. Mice were euthanized when tumors reached 2 cm in any dimension or at experimental endpoint (day 58), and tumors were harvested for analysis.

#### Drug Treatments

Doxycycline (Sigma, D9891) was dissolved in water and added to culture medium at a final concentration of 1 μg/mL. All-trans retinoic acid (ATRA; Thermo Fisher Scientific) was dissolved in DMSO and used at 1 μM. Arsenic trioxide (ATO; Sigma) was dissolved in 1 N NaOH and used at 1 μM. A485 (Selleckchem, S7393) was dissolved in DMSO and used at the indicated concentrations. Vehicle controls (DMSO or appropriate solvent) were included in all experiments.

### Genetic Engineering and Perturbations

#### CRISPR/Cas9 knock-in of EBF3-HiBiT-3×FLAG

To enable detection of endogenous EBF3 protein, we engineered NB4 cells to express EBF3 with a C-terminal HiBiT-3×FLAG tag using CRISPR/Cas9-mediated homology-directed repair (HDR). The sgRNA targeting the EBF3 C-terminus and the single-stranded oligodeoxynucleotide (ssODN) HDR donor template were designed using the IDT Alt-R HDR Design Tool (Integrated DNA Technologies). The HDR donor oligo contained the HiBiT-3×FLAG coding sequence flanked by homology arms and included phosphorothioate modifications at the terminal nucleotides to enhance stability.

Cas9 ribonucleoprotein (RNP) complexes were assembled by incubating Alt-R S.p. Cas9 Nuclease V3 (IDT) with Alt-R CRISPR-Cas9 sgRNA at room temperature for 10–20 minutes. NB4 cells were nucleofected with RNP complexes, HDR donor oligo, and Alt-R Cas9 Electroporation Enhancer (IDT) using the Lonza 4D-Nucleofector System with the SF Cell Line Kit and program CZ-100. Following nucleofection, cells were cultured in medium containing Alt-R HDR Enhancer V2 (1 μM; IDT) for 24 hours to promote HDR, then transferred to standard growth medium. Single-cell clones were isolated by limiting dilution, and successful knock-in was validated by Sanger sequencing and Inference of CRISPR Edits (ICE) analysis. The selected clone showed 100% knock-in efficiency at the targeted allele (Supplemental Fig. 1B-D). sgRNA and HDR donor oligo sequences are provided in Supplemental Table S7.

#### Lentiviral production and shRNA knockdown

Lentivirus was produced in HEK293T cells via transfection of psPAX2 (Addgene #12260), VSV-G (Addgene #8454), and lentiviral transfer plasmids using TransIT-X2 (Mirus). Viral supernatants were collected 48-72 hours post-transfection and filtered through 0.45-μm PVDF filters (Millipore). NB4 or NB4-R2 cells were transduced in the presence of 5 μg/mL polybrene (Sigma) with lentiviral particles from doxycycline-inducible shRNAs (tet-pLKO-puro, Addgene #21915) targeting EBF3, PML, a non-targeting scrambled control (SCR), or empty vector (EV). Transduced cells were cultured in tetracycline-free media and selected with 1 μg/mL puromycin (Thermo). shRNA sequences are provided in Supplemental Table S7.

#### Overexpression Constructs

For EBF3 overexpression studies, the human *EBF3* coding sequence (UniProt Q9H4W6-2) was cloned into the pSIN4-EF1α-IRES-Puro backbone (Addgene #61065) with an N-terminal FLAG tag. Patient-derived (C198F) and HADDS-associated (R163P, P177L) mutations were introduced by site-directed mutagenesis using the Q5 Site-Directed Mutagenesis Kit (NEB). An *eGFP-FLAG* construct in the same backbone served as a negative control. All constructs were verified by Sanger sequencing. Lentiviral production and transduction were performed as described above.

#### Antisense Oligonucleotide (ASO) Knockdown

ASOs were designed to target *EBF3* and synthesized by Integrated DNA Technologies (IDT). ASOs contained phosphorothioate backbone modifications throughout and 2′-O-methoxyethyl (2′- MOE) modifications at the five terminal nucleotides of both 5′ and 3′ ends to enhance stability and reduce toxicity. Primary APL cells (5 × 10⁴) were suspended in 20 μL SF Cell 4D-Nucleofector X Solution (Lonza) and mixed with 500 pmol ASO. This ASO amount enabled near-complete delivery to the cellular milieu. Nucleofection was performed using the CZ-100 program on the 4D- Nucleofector X Unit (Lonza). Cells were transferred to fresh primary culture medium and incubated at 37°C for subsequent experiments. ASO sequences:

### Target Sequence (5′→3′)

EBF3 mAmTmAmGmTGGATGCCGTTmGmTmTmGmG Negative Control mCmCmTmAmTAGGACTATCCmAmGmGmAmA N = deoxynucleotide; mN = 2′-O-methoxyethyl deoxynucleotide.

#### CRISPR Tiling Screen

To identify regulatory elements required for *EBF3* expression and cell fitness, we performed a CRISPR/Cas9 tiling screen across the *EBF3* locus (chr10:129809291-130018162, GRCh38). sgRNA sequences and efficiency scores were obtained from the UCSC Genome Browser CRISPR Targets track (Haeussler et al., 2016), which provides pre-computed Doench/Fusi efficiency scores for all NGG PAM sites in the human genome. The library comprised 1,999 sgRNAs targeting: (1) three intronic enhancer regions corresponding to PML::RARA-bound sites (E1: chr10:129883600-129884100, 52 sgRNAs; E2: chr10:129881500-129882100, 86 sgRNAs; E3: chr10:129870500-129872000, 156 sgRNAs; score threshold > 50); (2) the TSS region (chr10:129962000-129966000, 237 sgRNAs; score > 80); (3) the promoter region (chr10:129968000-129973000, 364 sgRNAs; score > 80); (4) intragenic tiling sgRNAs spanning the locus (968 sgRNAs; score > 90); (5) positive control sgRNAs targeting EBF3, PML, and RARA (4 sgRNAs each, from the Brunello library; Doench et al., 2016); and (6) non-targeting control sgRNAs (124 sgRNAs, randomly selected from the Brunello library). The pooled oligonucleotide library was synthesized by Twist Bioscience and cloned into lentiGuide-Puro (Addgene #52963) by Gibson assembly. Library representation was verified by amplicon sequencing prior to screening. sgRNA sequences and genomic coordinates are provided in Supplemental Table S7. NB4 cells were transduced with lentiCas9-Blast (Addgene #52962) and selected with blasticidin to generate NB4-Cas9 cells. NB4-Cas9 cells and parental NB4 cells (Cas9-negative control) were transduced with the pooled sgRNA library at low MOI (∼0.3), selected with puromycin (1 μg/mL), and maintained at ≥500× library representation (n = 3 biological replicates per condition). Cells were harvested at day 21 post-transduction. Genomic DNA was extracted using the Monarch Spin gDNA Extraction Kit (NEB), sgRNA cassettes were amplified by PCR with barcoded primers (Joung et al. 2017), and libraries were sequenced on an Illumina MiSeq. sgRNA read counts were quantified using MAGeCK (Li et al., 2014). Differential sgRNA abundance between NB4-Cas9 and NB4 conditions was determined using DESeq2, with size factors estimated from non-targeting control sgRNAs. Log2 fold change shrinkage was applied using the apeglm method. Negative log2 fold change (sgRNA depletion in Cas9+ cells) indicates targeting of sequences required for cell fitness. Feature-level comparisons were performed using Wilcoxon rank-sum tests.

#### CRISPRi Validation

For CRISPRi-mediated repression of EBF3 regulatory elements, NB4-EBF3-HiBiT cells were transduced with lentivirus encoding dCas9-KRAB (Lenti-dCas9-KRAB-blast, Addgene #89567). Individual sgRNAs targeting the *EBF3* TSS or intronic enhancer elements were cloned into LentiGuide-puro-NLS-GFP (Addgene #185473) and delivered by lentiviral transduction. sgRNA sequences are provided in Supplemental Table S7. At 72 hours post-transduction, 5,000 eGFP+ cells were sorted into individual wells by fluorescence-activated cell sorting (FACS) on a CytoFLEX SRT (Beckman Coulter), and EBF3 protein levels were measured by HiBiT lytic assay.

### Biochemical and Cellular Assays

#### Protein Extraction and Western Blotting

For whole-cell lysates, cells were resuspended in RIPA buffer (Boston BioProducts) supplemented with 1× EDTA-free protease/phosphatase inhibitor cocktail (Thermo Fisher Scientific). Lysates were sonicated using a Sonic Dismembrator 100 (Fisher Scientific) at setting 3 with 3-4 pulses of 15 seconds on/off on ice, followed by incubation for 1 hour at 4°C with end-over-end rotation. Lysates were clarified by centrifugation (15,000 × g, 20 min, 4°C), and protein concentrations were determined by BCA assay (Thermo Fisher Scientific).

Nuclear and cytoplasmic protein fractions were prepared using the NE-PER Nuclear and Cytoplasmic Extraction Reagents (Thermo Fisher Scientific, 78833) according to the manufacturer’s instructions, with protease inhibitors (Halt Protease Inhibitor Cocktail; Thermo Fisher Scientific) added to extraction buffers. Fraction purity was validated by Western blotting for GAPDH (cytoplasmic) and Lamin B1 (nuclear).

Proteins were separated on NuPAGE 4-12% Bis-Tris gels (Thermo Fisher Scientific) and transferred to 0.45-μm nitrocellulose membranes (Bio-Rad). Membranes were blocked with 3% non-fat milk in PBS-T (PBS with 0.1% Tween-20) and incubated overnight at 4°C with primary antibodies. After washing, membranes were incubated with HRP-conjugated secondary antibodies for 1 hour at room temperature. Chemiluminescence was detected using SuperSignal West Femto Maximum Sensitivity Substrate (Thermo Fisher Scientific) and imaged on an Amersham Imager 600 (GE Healthcare). Membranes were stripped with Restore PLUS Stripping Buffer (Thermo Fisher Scientific) for reprobing. HiBiT-tagged proteins were detected using the Nano-Glo HiBiT Blotting System (Promega). Images were processed with Fiji/ImageJ v2.3.0.

Primary antibodies: anti-FLAG M2 (Sigma-Aldrich, F1804; 1:2,000), anti-PML (Bethyl, A301-167A; 1:1,000), anti-RARA (CST, 1:1,000), anti-H3K27ac (CST, 8173S; 1:1,000), anti-ACTB (Sigma, A5316; 1:5,000), anti-Lamin B1 (Abcam, ab16048; 1:1,000), anti-GAPDH (CST, 2118S; 1:1,000). Secondary antibodies: anti-rabbit IgG HRP (GE Healthcare, NA9340; 1:5,000), anti-mouse IgG HRP (GE Healthcare, NA931; 1:5,000).

#### HiBiT Lytic Assay

Endogenous EBF3 protein levels were quantified using the Nano-Glo HiBiT Lytic Detection System (Promega). NB4-EBF3-HiBiT cells were plated in 96-well plates, and at the indicated timepoints, Nano-Glo HiBiT Lytic Reagent was added at a 1:1 volume. After 10 minutes of incubation at room temperature, luminescence was measured on a GloMax plate reader. For cell count normalization in A485 treatment experiments, parallel wells were assayed for cell viability using CellTiter-Glo (Promega).

#### Cell Viability Assays

Cell viability was measured using CellTiter-Glo Luminescent Cell Viability Assay (Promega). For shRNA knockdown assays, NB4 or NB4-R2 cells (1 × 10^4^ per well) were plated in 96-well plates with or without 1 μg/mL doxycycline. At the indicated timepoints (days 0, 3, and 6), an equal volume of CellTiter-Glo reagent was added to each well, incubated for 10 minutes at room temperature, and luminescence was measured on a GloMax (Promega) plate reader. Luminescence values were normalized to corresponding negative control for each condition.

For A485 dose-response experiments, NB4 or NB4-R2 cells were treated with the indicated concentrations of A485 or DMSO vehicle. Viability was measured at the indicated time points by CellTiter-Glo. IC50 values were calculated by fitting a four-parameter log-logistic model using the drc package in R.

#### Colony Formation Assay

NB4 cells transduced with doxycycline-inducible shRNA constructs were resuspended in IMDM supplemented with 2% FBS (StemCell Technologies) and plated in MethoCult (StemCell Technologies, H4230) at 1,000 cells per well in triplicate (6-well plates) with or without 1 μg/mL doxycycline. Colonies were counted after 14 days.

#### Competition Assay

NB4 cells stably expressing FLAG-EBF3 variants (WT, C198F, R163P, P177L) were mixed 1:1 with eGFP-FLAG control as a reference population. Mixed populations were cultured for 14 days, with the eGFP+ percentage measured via flow cytometry at the indicated timepoints. Flow cytometry was performed on a CytoFLEX LX (Beckman Coulter) and analyzed using CytExpert software.

#### RNA Isolation and RT-qPCR

Cells were harvested and total RNA was extracted using TRIzol Reagent (Thermo Fisher Scientific, 15596026) according to the manufacturer’s instructions. RNA was quantified by NanoDrop (DeNovix DS-11). cDNA was synthesized from 1 μg total RNA using the SuperScript IV First-Strand Synthesis System (Thermo Fisher Scientific, 18091050). Quantitative PCR was performed in triplicate using LightCycler 480 SYBR Green I Master (Roche, 04887352001) on a LightCycler 480 Instrument II. Gene expression was normalized to ACTB using the 2^−ΔΔCt method. Primer sequences are provided in Supplemental Table S7.

#### iDAPT-MS

Chromatin accessibility and associated protein microenvironments were profiled in NB4 cells using iDAPT (integrative DNA And Protein Tagging), as previously described (Lee et al., 2021). iDAPT employs a bifunctional Tn5 transposase-APEX2 peroxidase fusion protein (TP) that exploits the stable association of Tn5 with DNA after tagmentation to anchor APEX2 at sites of accessible chromatin. Upon addition of biotin-phenol and hydrogen peroxide, APEX2 catalyzes the formation of biotin-phenoxyl radicals that covalently label proximal proteins within a ∼20 nm radius, enabling identification of the protein microenvironment at open chromatin regions.

NB4 cells were treated with 1 μM ATRA or vehicle (DMSO) for 48 hours. Nuclei were isolated by resuspension in hypotonic lysis buffer supplemented with 0.1% NP-40, and 500,000 nuclei per replicate were tagmented with the TP fusion protein or negative controls (Tn5 alone or APEX2 alone). For NB4 cells, an endogenous peroxidase blocking step (0.03% H₂O₂, 0.1% NaN₃, 30 min on ice) was performed after nuclear extraction and before tagmentation. Tagmentation was performed at 37°C for 30 minutes, after which unbound protein was removed by washing. Proximity labeling was initiated by addition of 500 μM biotin-phenol and 1 mM H₂O₂ for 1 minute at room temperature, then quenched with sodium ascorbate, Trolox, and sodium azide. Labeled nuclei were lysed in RIPA buffer supplemented with protease inhibitors, sonicated, and treated with benzonase to degrade nucleic acids. Clarified lysates were quantified by detergent- compatible Bradford assay, and biotinylated proteins were enriched by streptavidin pulldown. Negative controls establish the specificity of the approach: Tn5 alone binds open chromatin but lacks peroxidase activity, controlling for endogenous biotinylation and nonspecific protein capture; APEX2 alone possesses peroxidase activity but is not retained at chromatin after washing, controlling for untethered labeling. Robust biotinylation requires the intact fusion protein together with both biotin-phenol and H₂O₂.

Enriched proteins were digested on-bead with trypsin and labeled with TMT 10-plex reagents. Pooled TMT-labeled peptides were fractionated by basic pH reversed-phase HPLC into 96 fractions consolidated into 24 super-fractions, desalted via StageTip, and analyzed by LC– MS/MS on an Orbitrap Fusion mass spectrometer coupled to a Proxeon EASY-nLC 1200 using an SPS-MS3 method to minimize reporter ion interference. Mass spectra were searched against the human UniProt database using a Sequest-based pipeline with TMT tags on lysine residues and peptide N-termini (+229.163 Da), carbamidomethylation of cysteine (+57.021 Da), and oxidation of methionine (+15.995 Da) as modifications. Peptide-spectrum matches were filtered to 1% FDR by linear discriminant analysis. For quantitation, summed signal-to-noise ratios were extracted for each TMT channel, excluding PSMs with reporter summed signal-to-noise < 100 or isolation specificity < 0.7. PSM intensities were normalized using the median intensity of streptavidin and trypsin PSMs per sample as internal standards, log₂-transformed, and collapsed to proteins by arithmetic average. Differential protein abundances were determined using the limma package in R.

For DMSO versus ATRA comparisons, n = 3 biological replicates per condition were analyzed; for TP versus negative control comparisons, n = 3 (TP) and n = 4 (controls) replicates were used. The TP and MEDS adaptor preparations were performed as previously described (Lee et al., 2021). Data from this previously published dataset (Lee et al., 2021) are reanalyzed and visualized in Fig. 1C and Supplemental Fig. 1A.

#### RNA-seq Library Preparation and Sequencing

Total RNA was extracted using TRIzol and quantified by NanoDrop and TapeStation 4200 (RNA ScreenTape; Agilent) or 2100 Bioanalyzer (Agilent). For NB4 shRNA libraries, total RNA-seq libraries were prepared from 500 ng total RNA using the KAPA mRNA HyperPrep Kit (Roche) with RiboErase (HMR) according to the manufacturer’s instructions. Library quality was assessed by TapeStation (High Sensitivity D1000 ScreenTape; Agilent), and libraries were pooled and sequenced on a NovaSeq 6000 (Illumina). For NB4 overexpression and ATRA treatment libraries, total RNA was submitted to Novogene for library preparation and sequencing. Poly(A)-selected mRNA libraries were prepared and sequenced on a NovaSeq 6000 (Illumina).

#### ATAC-seq

ATAC-seq was performed using the Omni-ATAC protocol with modifications as previously described (Corces et al., 2017; Lee et al., 2021). Briefly, 100,000 cells were lysed in lysis buffer (10 mM Tris-HCl pH 7.5, 10 mM NaCl, 3 mM MgCl₂, 0.01% digitonin, 0.1% Tween-20, 0.1% NP- 40) and nuclei were pelleted. Tagmentation was performed with Tn5 transposase loaded with MEDS-A/B adapters in tagmentation buffer (20% dimethylformamide, 10 mM MgCl₂, 20 mM Tris-HCl pH 7.5, 33% PBS, 0.01% digitonin, 0.1% Tween-20) for 30 min at 37°C with thermomixing. DNA was purified using DNA Clean & Concentrator-5 (Zymo Research) and amplified using barcoded primers. Optimal PCR cycle number was determined by qPCR. Libraries were analyzed by TapeStation (High Sensitivity D1000) and sequenced on a NovaSeq 6000.

#### CUT&RUN

pAG-MNase was purified from Rosetta2 cells as described (Lee et al., 2021). CUT&RUN was performed as previously described (Skene and Henikoff, 2017) with modifications. Briefly, 500,000 NB4 cells per reaction were washed in wash buffer (20 mM HEPES pH 7.5, 150 mM NaCl, 0.5 mM spermidine, 1× protease inhibitor cocktail) and bound to Concanavalin A-coated magnetic beads. Cells were incubated overnight at 4°C with primary antibodies (anti-FLAG, CST 14793S, 1:50; anti-FLAG M2, Sigma F1804, 1:100) in dig-wash buffer (wash buffer with 0.01% digitonin). After washing, pAG-MNase (2 μg/mL) was added and incubated for 1 hour at 4°C. MNase cleavage was activated by addition of CaCl₂ (2 mM final) at 0°C for 1 hour. Reactions were stopped, DNA was purified by phenol-chloroform extraction, and libraries were prepared using the NEBNext Ultra II DNA Library Prep Kit (NEB). Libraries were sequenced on a NovaSeq 6000.

#### CUT&Tag

CUT&Tag was performed using a modified protocol (Kaya-Okur et al., 2019). Primary APL cells (5 × 10⁵ per reaction) were bound to Concanavalin A beads in a 1:10 ratio of binding buffer (20 mM HEPES pH 7.5, 10 mM KCl, 1 mM CaCl₂, 1 mM MnCl₂) to wash buffer (20 mM HEPES pH 7.5, 150 mM NaCl, 0.5 mM spermidine, 1× protease inhibitor cocktail). Bead-bound cells were incubated overnight at 4°C with primary antibodies in antibody buffer (wash buffer with 0.05% digitonin, 2 mM EDTA and 0.1% BSA). Antibodies used: anti-RNA polymerase II (clone CTD4H8; Millipore Sigma, 05-623-25UG; 1:50). After washing, cells were incubated with secondary antibody (rabbit anti-mouse IgG; abcam, ab46540; 1:100) in dig-wash buffer (wash buffer with 0.05% digitonin) for 30–60 min. pA-Tn5 transposome was added in dig-300 buffer (20 mM HEPES pH 7.5, 300 mM NaCl, 0.5 mM spermidine, 0.01% digitonin, 1× protease inhibitor cocktail) for 1 hour. Tagmentation was performed (dig-300 with 10 mM MgCl₂) for 1 hour at 37°C. DNA was purified with Zymo Research DNA Clean & Concentrator-5 columns, and libraries were prepared using NEBNext reagents. Libraries were sequenced on a NovaSeq 6000.

### Bioinformatic Analysis

#### RNA-seq Analysis

For differential expression analysis, raw sequencing reads were pseudo-aligned to the human transcriptome (Ensembl v104, GRCh38) using Salmon v1.10.3 quant with options “--seqBias -- useVBOpt --gcBias --posBias --numBootstraps 30”. Transcript-level abundances were summarized to gene level using tximport v1.38.1. Differential expression analysis was performed using DESeq2 v1.50.2 in R. For visualization, count data were transformed using the variance-stabilizing transformation (VST). Principal component analysis and t-SNE dimensionality reduction (Rtsne v0.17) were performed on VST-transformed counts. For APL patient sample clustering analyses, only genes with significant differential expression upon EBF3 knockdown (FDR < 0.05, |log₂ fold change| > 1; 1,559 genes) were used as features.

For genome track visualization, reads were aligned to the human genome (Ensembl v104, GRCh38) with STAR v2.5.2b. Duplicates were removed using Picard v2.8.0. Read density was counted over a 150 bp window sliding 20 bp across the genome and then converted to a bigwig file using ucsc-tools v475. Genome tracks were visualized using a custom script in R.

Receiver operating characteristic (ROC) curves were generated to assess the ability of EBF3 expression to classify APL versus non-APL AML samples in R.

#### ATAC-seq and CUT&Tag Analysis

Paired-end reads were trimmed with TrimGalore v0.4.5 to remove adapter sequences. Reads were aligned to the human genome (GRCh38) using Bowtie2 v2.5.4 with options “--no-unal --no-discordant --no-mixed -X 2000”. Mitochondrial reads were removed, and duplicates were marked using Picard v2.8.0. For visualization, Tn5 insertion sites were adjusted (+4 bp for forward strand, −5 bp for reverse strand), and bigwig files were generated and visualized as described above. Differential accessibility analysis was performed using DESeq2 on read counts within CUT&RUN peak regions.

#### CUT&RUN Analysis

Paired-end reads were trimmed with TrimGalore and aligned to GRCh38 using Bowtie2 with options “--no-unal --no-discordant --no-mixed --dovetail -I 25 -X 700”. Duplicates were removed with Picard. EBF3 CUT&RUN peaks were called using MACS2 with options “callpeak -q 0.01 -- keep-dup all”. Peak reproducibility was assessed using IDR, and peaks with IDR < 0.05 were retained. Motif enrichment analysis was performed using HOMER v4.9 with motifs from JASPAR 2024 (Rauluseviciute et al., 2024). Target genes were assigned to peaks within 50 kb of transcription start sites.

#### Gene Set Enrichment Analysis

Gene set enrichment analysis (GSEA) was performed using fgsea in R with genes ranked by log₂ fold change and excluding those with FDR > 0.5. The MSigDB Hallmark gene set collection (Liberzon et al., 2015) was used for pathway analysis.

#### Mutation and Variant Allele Frequency Analysis

For the APL2012 cohort, DNA-based variant allele frequencies (VAF) were obtained from reported VAFs from DNA sequencing (Lin et al., 2021). RNA-based VAFs were recalculated from RNA-seq reads at variant positions using a GATK pipeline.

### Statistical Analysis

Statistical analyses were performed in R. Two-tailed tests were used unless otherwise specified. For comparisons between two groups, Welch’s t-test (for normally distributed data) or Wilcoxon rank-sum test (for non-parametric data) was used. For comparisons involving multiple groups, one-way ANOVA with post-hoc Dunnett’s test (comparison to control) or pairwise comparisons with Holm-Sidak correction were used. For survival analysis, log-rank (Mantel-Cox) test was used. Sample sizes are indicated in figure legends. No statistical methods were used to predetermine sample size. Experiments were not randomized, and investigators were not blinded to group allocation during experiments or outcome assessment.

### AI/LLM Usage

Artificial intelligence tools (Claude, Anthropic) were used to assist with manuscript preparation, including editing of text, and to assist with code generation for bioinformatics analyses. All AI-generated text was reviewed, verified, and revised by the authors. All AI-generated code was reviewed, tested, and validated by the authors against expected outputs. No AI tools were used in experimental design or data collection.

